# Lipids Are Involved in Heterochromatin Condensation: A Quantitative Raman and Brillouin Microscopy Study

**DOI:** 10.1101/2025.02.26.640449

**Authors:** Masato Machida, Shinji Kajimoto, Ren Shibuya, Mayu Isono, Mai Watabe, Yukako Oma, Kayo Hibino, Kentaro Fujii, Masaki Okumura, Masahiko Harata, Atsushi Shibata, Takakazu Nakabayashi

**Author notes:** **Corresponding authors:** Shinji Kajimoto; Takakazu Nakabayashi.

## Abstract

Chromatin, a fundamental component of eukaryotic genomes, is categorized into euchromatin and heterochromatin, which play distinct roles in gene regulation. Although these two chromatin states are distinguished by their degree of condensation, quantitatively measuring the degree of chromatin condensation, as well as the physical properties of chromatin in living cells, remains challenging. In this study, label-free *in situ* quantitative imaging was performed using a Raman-Brillouin microscope to visualize the spatial distribution of molecular concentration and viscoelasticity within the nuclear environment of a living cell. A quantitative concentration distribution image of each intracellular biomolecule was obtained by combining Raman imaging with multivariate curve resolution analysis, using a water Raman band as an internal standard. Simultaneous Raman-Brillouin imaging enables the quantitative visualization of viscoelasticity within a cell. Using this approach, we found that, in addition to DNA, heterochromatin is enriched in lipids and that lipids play a critical role in heterochromatin formation, determining its mechanical properties. These findings provide new insights into the mechanism of heterochromatin formation and its chemical and physical properties, leading to a comprehensive understanding of gene regulation and nuclear organization.

## Introduction

Chromatin, the foundation of eukaryotic genome organization, plays a central role in regulating essential cellular functions such as gene expression and DNA repair^1,2,3,4^. Proper DNA organization in the chromatin is critical for cell survival, differentiation, and development. Abnormalities in chromatin structure or chemical composition can lead to various diseases, including cancer and neurodegenerative diseases^5,6,7^. Dynamic changes in the chromatin architecture affect transcriptional regulation, cell cycle progression, and genome stability. Despite its importance in biology ^8^, ^9^, chromatin organization and regulation remain poorly understood.

A key challenge in chromatin biology is quantifying the chemical composition. Based on the degree of condensation, chromatin can be categorized into two distinct types: euchromatin and heterochromatin. Euchromatin, characterized by a loosely packed structure, is highly transcriptionally active, whereas heterochromatin is densely packed and transcriptionally inert^10,11^. Heterochromatin can be visualized using DNA-specific staining dyes DAPI or Hoechst, appearing as spots several times brighter than the surrounding nucleoplasm. These observations support the idea that the DNA concentration in heterochromatin is several times higher than that in euchromatin which occupies the nucleoplasm^12,13,14^. However, the precise concentrations of DNA and other chemical components within heterochromatin and euchromatin remain elusive, and consequently, the degree of condensation in densely packed heterochromatin cannot be quantitatively evaluated. The lack of quantitative discussion is partly attributed to fluorescence microscopic measurements, which provide absolute fluorescence intensity and are inherently non-quantitative. Refractive index measurements also suggest that heterochromatin is more condensed^12, 15^; however, they do not provide molecular information, leaving it unclear what is enriched and to what extent.

Understanding the mechanical properties of chromatin is equally important^16,17,18,19,20^. These properties directly influence the accessibility of transcription factors to genes that regulate gene expression^21,22^,^23^. Certain genetic mutations and pathological conditions, such as cancer, are also known to alter chromatin architecture and its mechanical properties^24,25^. Abnormal chromatin organization in cancer cells leads to aberrant gene expression^26,27^. Understanding chromatin mechanics could provide new therapeutic and diagnostic approaches; however, chromatin exhibits different viscoelastic behaviors depending on the observation method. Chromatin behaves as a liquid when nanoscale dynamics are observed using single-molecule fluorescence imaging ^28^. In contrast, chromatin behaved as a solid when mesoscale dynamics were examined using fluorescence recovery after photobleaching ^29^. A quantitative method is required to evaluate the viscoelasticity of chromatin.

Few methods have been proposed to measure molecular concentrations in a single living cell. Optical tomography visualizes the three-dimensional distribution of molecular density in a living cell by assuming that molecular density is proportional to the refractive index, regardless of the molecular composition^30^. Oh et al. proposed normalized Raman imaging using stimulated Raman microscopy, in which Raman signals in the C–H and O–H stretching band regions were decomposed into lipids, proteins, and water^31^. In the present study, we introduce simultaneous Raman and Brillouin imaging to quantitatively visualize molecular concentrations and physical properties of heterochromatin and euchromatin within a single living cell. In previous reports^32,33^, we demonstrated that, by utilizing the water Raman band in the extracellular regions as an internal standard, local concentrations of nucleic acids and drug molecules delivered can be determined based on Raman imaging of a living cell. Here, we applied multivariate curve resolution-alternating least squares (MCR-ALS) to the Raman image of a single cell to distinguish and quantify concentrations of DNA, RNA, proteins, lipids, and water^34,35^. A Brillouin image was obtained simultaneously with the Raman image, allowing visualization of the distribution of viscoelastic properties in the living cell. We simultaneously quantified the chemical composition, concentration, and stiffness of heterochromatin and euchromatin in living cells. Moreover, we demonstrated that lipids are involved in heterochromatin formation and play a pivotal role in the condensation mechanism.

## Results

### Conventional Raman imaging allows label-free visualization of heterochromatin in living NIH3T3 cells in interphase

The typical Raman images and Raman spectra of living NIH3T3 cells are presented in Fig. 1. In NIH3T3 cells, heterochromatin was particularly condensed and observed as chromocenters (Supplementary Fig. 1)^36,37^. After obtaining the Raman images using a Raman microscope, the observed cells were fixed and stained with a DNA-staining dye, DAPI, on the microscope stage to visualize heterochromatin in the same cell (Fig. 1b). DAPI fluorescence revealed the presence of several densely stained regions in the nucleus, identifying DNA-packed heterochromatin as chromocenters. Because the intensity of each Raman band is proportional to the local concentration of the corresponding molecule, Raman images obtained using different Raman bands reflected the concentration distribution of specific biomolecules, that is, nucleic acids (788 cm^-1^, Fig. 1c), proteins (1003 cm^-1^, Fig. 1d), and lipids (2850 cm^-1^, Fig. 1e). The assignments of Raman bands are summarized in Supplementary Table 1. Most biomolecules contain C–H bonds; therefore, the Raman image obtained with the C–H stretching band reflects the local concentration of all biomolecules (Fig. 1f). The C–H Raman image exhibits a lower intensity in the nucleus than in the surrounding cytoplasm, revealing that the nucleus is less crowded with biomolecules than the surrounding cytoplasmic regions, which is consistent with our previous reports^38,39,40^. Strong C–H Raman intensities were observed in several regions within the nucleus, some of which corresponded to nucleoli observed in the bright-field image (Fig. 1a). The remaining strong regions overlapped with the chromocenters identified in the DAPI image (Fig. 1b), indicating that the heterochromatin was more densely packed with biomolecules than the nucleoplasm.

**Fig. 1.**
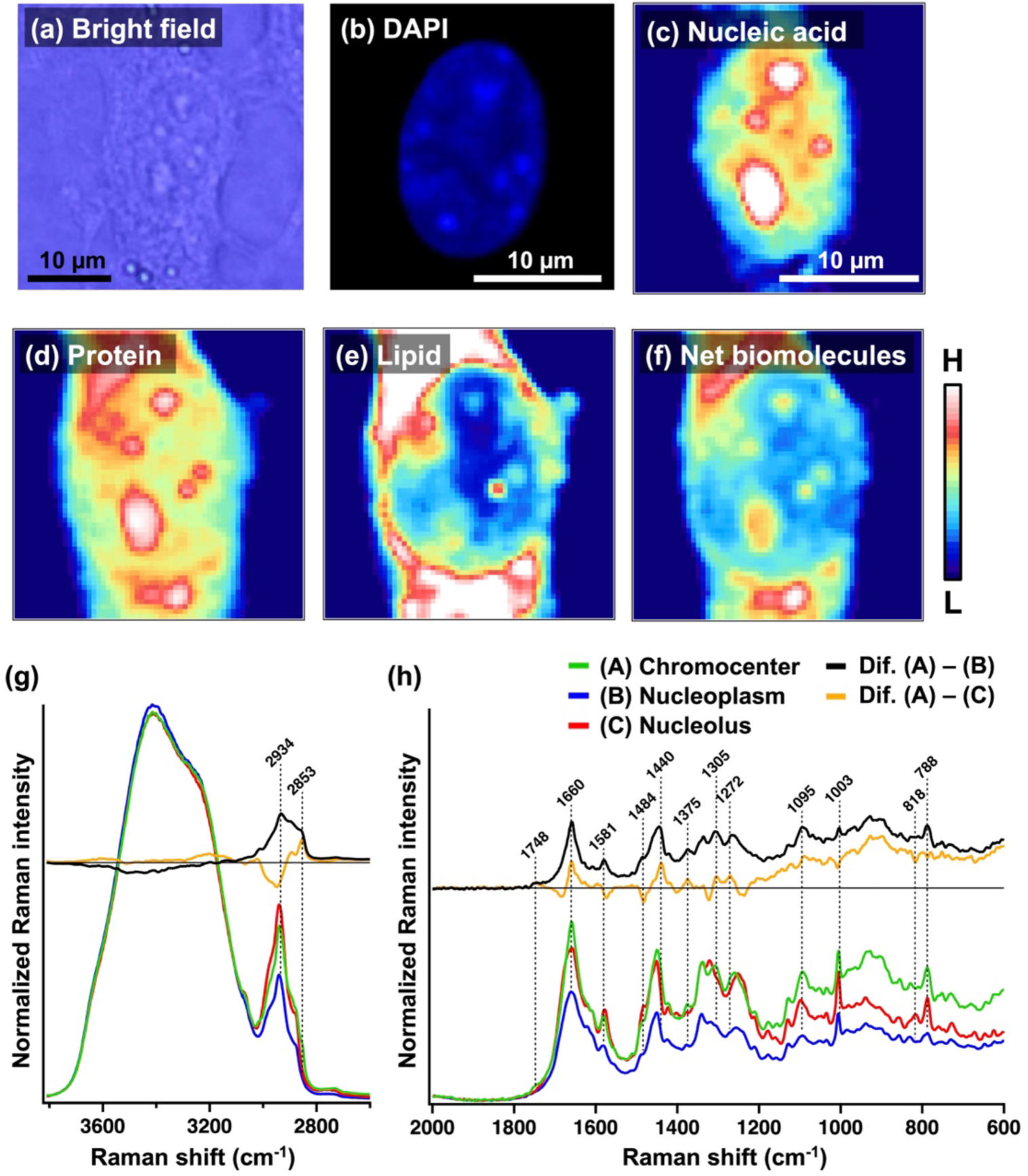
Raman images and spectra of a living NIH3T3 cell. (a-g) Bright-field (a), fluorescence (b), and Raman (c-f) images of an NIH3T3 cell. The fluorescence image was obtained after obtaining the Raman images, followed by cell fixation and staining with DAPI. Raman images were obtained by mapping the intensity of the pyrimidine band (c, 770-800 cm^-1^), phenylalanine band (d, 995-1020 cm^-1^), the lower region of the C–H stretching band (e, 2840-2860 cm^-1^) and the entire C–H stretching band (f, 2800-3030 cm^-1^). (g, h) Raman spectra of the chromocenter (green), nucleoplasm (blue), and nucleolus (red) of NIH3T3 cells in the higher wavenumber region (g) and the fingerprint region (h). Difference Raman spectra were obtained by subtracting the spectra of the nucleolus (yellow) and nucleoplasm (black) from the spectrum of the chromocenter. *n* = 6 cells.

Raman images of nucleic acids (Fig. 1c) and proteins (Fig. 1d) show that nucleoli and heterochromatin are brighter than their surroundings, indicating that both nucleoli and heterochromatin contain higher concentrations of proteins and nucleic acids than the nucleoplasm. Although distinguishing between nucleoli and heterochromatin in these Raman images was difficult, heterochromatin was identified as bright areas in the Raman image of lipids (Fig. 1e), implying that heterochromatin was lipid-rich compared with the surrounding nucleoplasm and nucleolus. This result was more apparent in the average Raman spectra of these regions and their Raman difference spectra (Fig. 1g, h). Positive bands in the difference spectrum between the chromocenter and nucleolus (chromocenter – nucleolus) were mainly assigned to lipids: ester C=O at 1745 cm^-1^, C=C bond at 1657 cm^-1^, CH2 bending at 1437 cm^-1^, and choline head group at 720 cm^-1^, indicating that heterochromatin contains more lipids than the nucleolus and nucleoplasm.

Raman spectroscopy allowed a detailed discussion of nucleic acid distribution within the nucleus. Both DNA and RNA exhibit a Raman band at 788 cm^-1^, attributed to the pyrimidine ring, but only RNA shows a sub-peak around 818 cm^-1^ (Supplementary Fig. 2). Upon comparing Raman spectra of the chromocenter, nucleolus, and nucleoplasm, we confirmed that both the chromocenter and nucleolus had nucleic acid concentrations several times higher than those of the nucleoplasm, whereas the nucleolus was enriched with a high concentration of RNA, and chromocenters contained a higher abundance of DNA (Fig. 1h).

### MCR Raman imaging enables quantitative molecular imaging of a living cell using a water Raman band as an intensity standard

We applied MCR-ALS analysis to Raman images to quantitatively visualize the distribution of each biomolecule within a living cell. We analyzed a Raman image (Supplementary Fig. 3) containing 3,600 Raman spectra and obtained 10 spectral components (Supplementary Fig. 4), along with their corresponding distribution images (Fig. 2). A water density image (Fig. 2c) was obtained by mapping the integrated intensity of the O–H stretching band of water (3500–3600 cm^-1^). The Raman spectrum of water was subtracted from each intracellular Raman spectrum, and the Raman spectra of the fingerprint region were subjected to MCR-ALS analysis. Comparing MCR spectra with Raman spectra of solution of different biomolecules, we identified that the obtained MCR spectra were derived from proteins, lipids, DNA, and RNA (Supplementary Fig. 5). The remaining MCR spectra were assigned to the resonance Raman spectra of cytochromes *b* and *c* and broad background emissions due to glass and biomolecular autofluorescence. The Raman spectrum of each cellular compartment was reproduced using six main components: proteins, RNA, DNA, lipids, and cytochromes *c* and *b* (Supplementary Fig. 6).

**Fig. 2.**
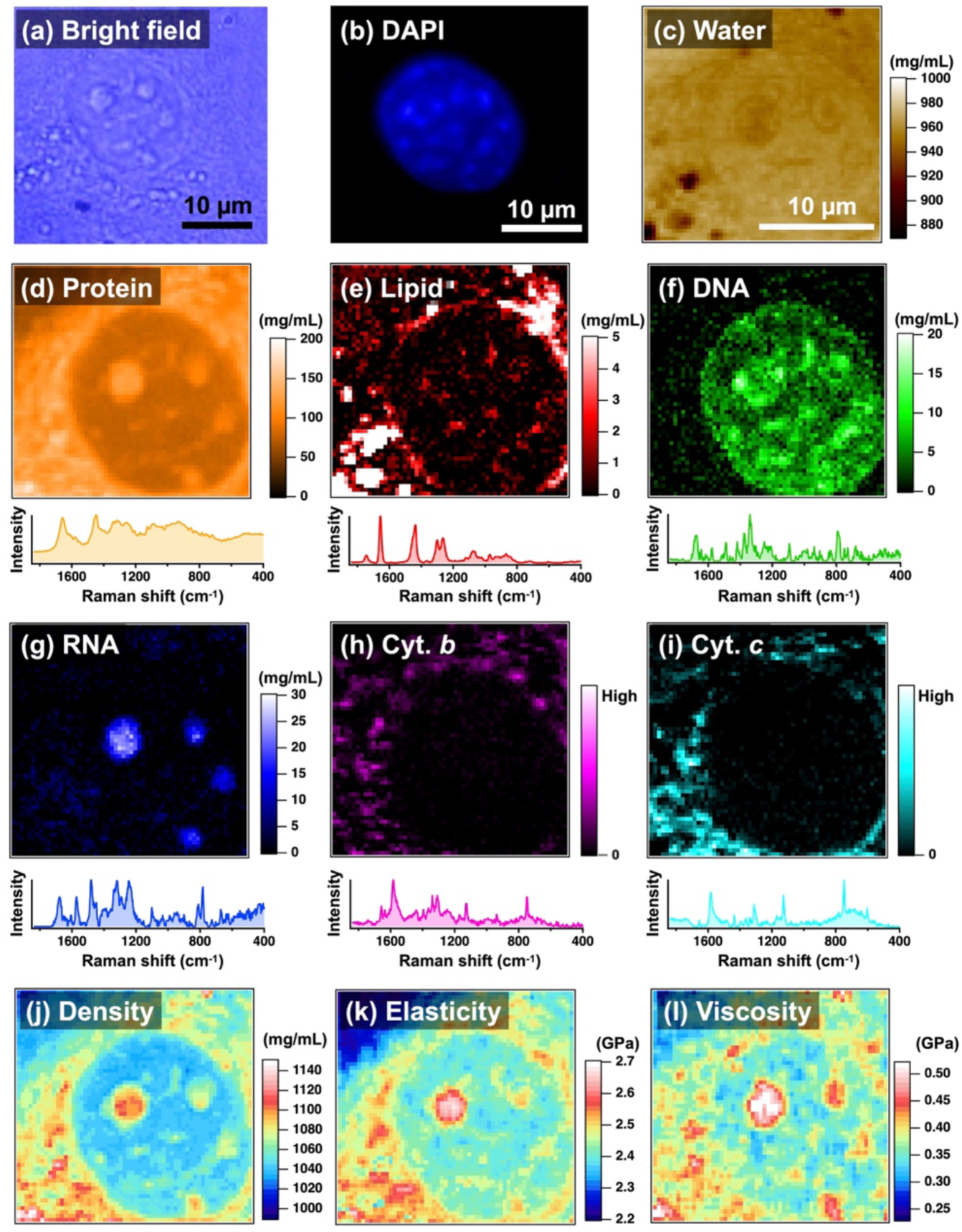
Biomolecular concentration and viscoelastic images of a living NIH3T3 cell. (a-c) Bright-field (a), fluorescence (b), and water Raman (c) images of an NIH3T3 cell. The water Raman image was obtained by mapping the intensity of the water Raman band relative to the water band in the extracellular region, in which the water density is 998 mg/mL, showing the absolute water density distribution intra- and extracellularly. (d-i) MCR images (top) and the corresponding MCR component spectra (bottom) assigned to proteins (d), lipids (e), DNA (f), RNA (g), cytochrome *b* (h), and cytochrome *c* (i). The intensity of each MCR image was converted to concentration using the calibration line of each biomolecule. (j) Density distribution image of the NIH3T3 cell. (k, l) Viscoelasticity images of the cell obtained using the density distribution, the corresponding Brillouin image, and the refractive index map in Supplementary Fig. 17.

The concentration was determined from the obtained MCR images using the Raman band of water as an internal standard. Raman scattering is inherently weak, and the measurement of absolute intensities often leads to considerable uncertainties in the quantitative analysis. However, given that the concentration of extracellular water remains constant, the Raman band of extracellular water can be used as an intensity standard to correct for experiment-to-experiment variations under optical conditions, thereby providing quantitative information on Raman intensities of cells. We first prepared calibration lines for proteins, lipids, DNA, and RNA by measuring Raman spectra of their solutions at different concentrations and plotting the relative intensity of a characteristic Raman band for each biomolecule to the O-H stretching band of water against the concentration (Supplementary Fig. 7-10). Subsequently, we substituted the intensity at each pixel of each MCR image relative to that of the O-H stretching band of extracellular water into the calibration line and obtained the concentration distribution of the biomolecule.

The color scale of MCR images in Fig. 2 does not represent the relative Raman band intensity but rather the absolute concentration obtained in “mg/mL”. Proteins, lipids, DNA, RNA, and water are the main components of living cells; therefore, the total density distribution within the cell was evaluated by a simple summation of these images (Fig. 2j). Upon examining the MCR images, we successfully visualized the distinct distributions of RNA and DNA: RNA was mainly concentrated in the nucleolus, whereas DNA was dispersed throughout the entire nuclear region. DNA is not homogeneously distributed in the nucleus, and there are several micrometer-sized spots with high DNA concentrations. These DNA-concentrated spots overlapped with the highly stained regions in the DAPI fluorescence image (Fig. 2b), indicating that the concentrated spots in the DNA image were chromocenters, that is, heterochromatin.

The average concentration of each biomolecule and the total density were evaluated for different nuclear regions based on the MCR images (Fig. 3 and Table 1). The DNA concentration in the heterochromatin was four times higher than that in the nucleoplasm, which is predominantly composed of euchromatin. This result is consistent with previous reports based on fluorescence and refractive index measurements^12,13,14^, indicating that heterochromatin is enriched in DNA by severalfold compared with euchromatin. In contrast, the differences in protein concentrations and total densities between heterochromatin and the nucleoplasm were relatively small (approximately 10%). In summary, the concentration of nucleosomes comprising DNA and histone proteins in heterochromatin was four times higher than that in euchromatin. Conversely, heterochromatin and euchromatin had nearly similar overall protein concentrations and total densities.

**Fig. 3.**
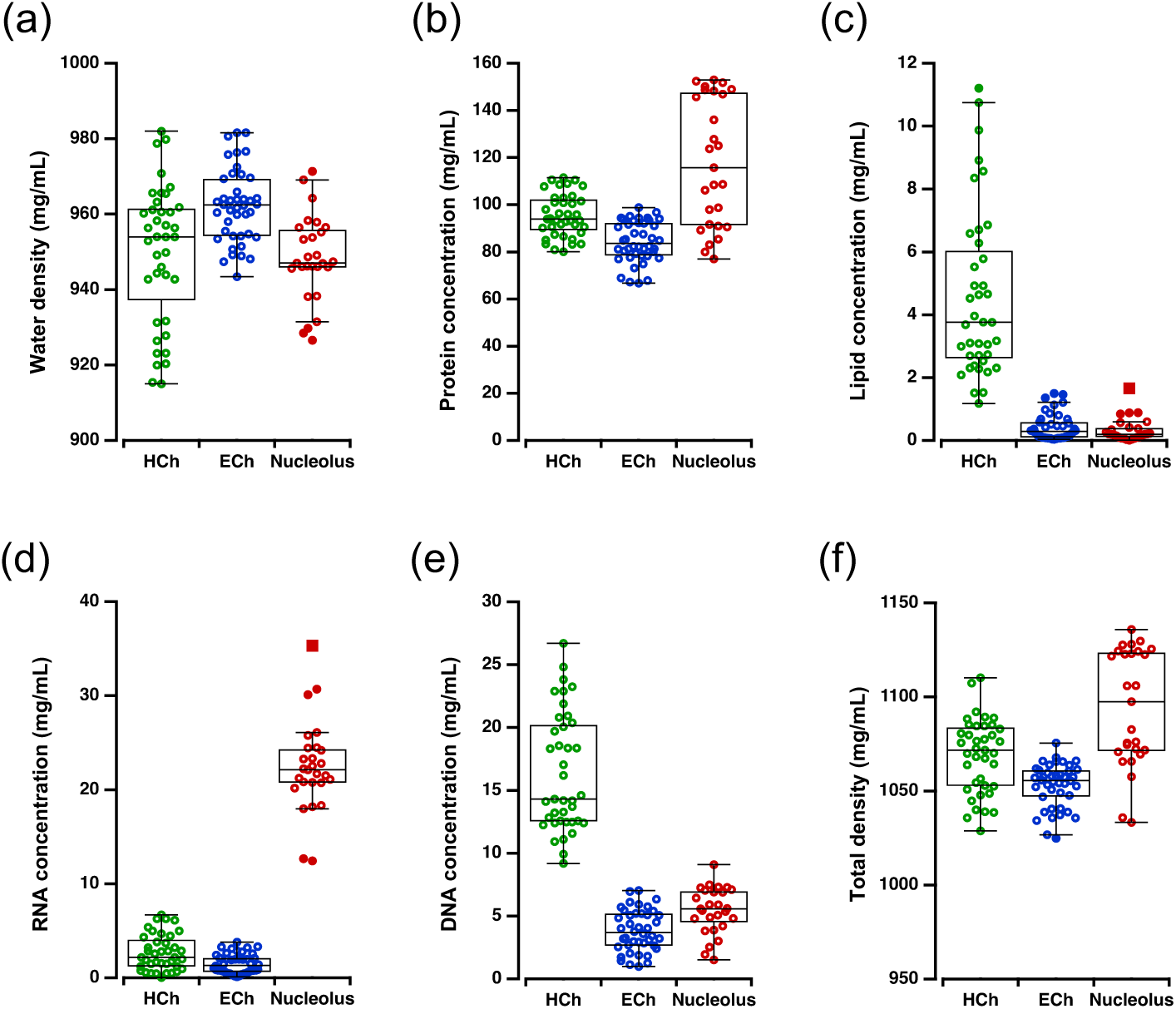
Concentration of biomolecules in each region of the nucleus. (a-f) The concentration of water (a), proteins (b), lipids (c), DNA (d), and proteins (e) and the total density (f) in heterochromatin (HCh), euchromatin (ECh), and the nucleolus. Closed circles and squares represent outer and far outer, respectively. *n* = 8 cells.

**Table 1.** Average biomolecular concentrations, refractive indices, and viscoelastic properties of heterochromatin, euchromatin, and nucleolus in living NIH3T3 cells. Concentrations of biomolecules and water were evaluated from the band intensity of each Raman band relative to the Raman band of extracellular water. The total density of each compartment was calculated as the summation of DNA, RNA, proteins, lipids, and water. The refractive indices were measured using a holographic microscope. The longitudinal and loss moduli of each compartment were estimated using the peak position and bandwidth of the Brillouin band and the density and refractive index of each compartment.

|  | Heterochromatin | Euchromatin | Nucleolus | Medium |
| --- | --- | --- | --- | --- |
| DNA conc. (mg/mL) | 16.4 | 3.8 | 5.4 | 0 |
| RNA conc. (mg/mL) | 2.7 | 1.5 | 22.4 | 0 |
| Protein conc. (mg/mL) | 95.5 | 84.3 | 117.8 | 0 |
| Lipids conc. (mg/mL) | 4.6 | 0.4 | 0.3 | 0 |
| Water conc. (mg/mL) | 948 | 962 | 950 | 998 |
| Density (mg/mL) | 1069 | 1053 | 1094 | 998 |
| Refractive index | 1.366 | 1.351 | 1.368 | 1.335 |
| Brillouin shift (GHz) | 7.90 | 7.83 | 8.10 | 7.42 |
| Longitudinal modulus (GPa) | 2.53 | 2.50 | 2.71 | 2.24 |
| FWHM (GHz) | 1.37 | 1.25 | 1.49 | 0.89 |
| Loss modulus (GPa) | 0.440 | 0.400 | 0.499 | 0.265 |

In the nuclear region, lipids were prominently observed only within the heterochromatin (Figs 2e and 3c). Merging images of DNA and lipids, we observed that lipid-rich regions overlap with DNA-rich heterochromatin (Supplementary Fig. 11). This result is consistent with the differences in Raman spectra (Fig. 1g, h). Overlapping DNA and lipid distribution was also observed in living HeLa cells, indicating that the lipid enrichment in heterochromatin was not specific to NIH3T3 cells (Supplementary Fig. 12, 13). To determine the chemical structures of lipids in heterochromatin, we extracted Raman spectra from the nuclear regions of multiple cells and applied MCR analysis to these spectra to obtain six-component spectra (Supplementary Fig. 14). The MCR spectrum assigned to lipids had a spectral profile closely resembling that of dioleoylphosphatidylcholine (DOPC), exhibiting a band at 719 cm^-1^ due to the choline group (Supplementary Fig. 15)^41,42^. The MCR spectrum, as well as the difference spectra in Fig. 1, exhibited an intensity ratio of approximately 1:1 between the C=C band at 1658 cm^-1^ and the CH2 scissors band at 1442 cm^-1^. Given that the intensity ratio *I*C=C/*I*CH2 is proportional to the unsaturation level of lipids (Supplementary Fig. 16)^43^, this result indicated that the average unsaturation level of lipids observed in heterochromatin was 1.4 times higher than that in DOPC, with an intensity ratio of 0.7. Note that when determining the biomolecular concentrations of heterochromatin, we analyzed only the internal heterochromatin, excluding peripheral heterochromatin (i.e., lamin-associated domains), to exclude the influence of the nuclear membrane adjacent to the peripheral heterochromatin.

### Simultaneous Raman and Brillouin imaging shows heterochromatin is harder than euchromatin

Simultaneous measurement of Brillouin images allowed the visualization of elasticity and viscosity distributions within a cell. Brillouin scattering is based on the interaction of acoustic phonons in materials with incident light, with estimated values reflecting the viscoelastic responses of materials in the GHz range^44,45,46,47^. Determining the viscoelastic properties of a sample from the Brillouin image requires obtaining the distributions of the refractive index and density within the cell. This is because the viscoelastic moduli obtained through Brillouin scattering are related to these parameters as follows^48,49^.

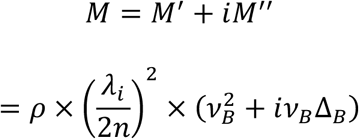

Where *M* represents the viscoelasticity, the real part *M*’ denotes the storage modulus (i.e., elasticity), and the imaginary part *M*’’ represents the loss modulus (i.e., viscosity). *ρ* is the total density, *λi* is the wavelength of the incident light (532 nm), *n* is the refractive index, *ϖB* is the peak position of the Brillouin band, and *11B* is the FWHM of the Brillouin band.

The refractive index of each cellular region of NIH3T3 cells was measured using a holographic microscope (Table 1, Supplementary Fig. 17). To obtain the refractive index distribution within the Raman/Brillouin-imaged cell, we segmented each Raman image into heterochromatin, nucleoli, nucleoplasm, lipid droplets, and other cytoplasmic regions based on the corresponding MCR image. We then constructed a refractive-index map of the cell using the average refractive index of each region (Supplementary Fig. 18). The variation in the refractive index in each nuclear region was less than 1%; therefore, the errors in the obtained refractive index map are expected to be ≤ 1%, at least in the nucleus. The density distribution within the cell was calculated as the sum of the densities of water and the biomolecules (Fig. 2j). Thus, we visualized the distribution of elasticity and viscosity within the cell by performing simultaneous Raman and Brillouin imaging using the above equations (Fig. 2k, l and Table 1).

Simultaneous Raman and Brillouin imaging enabled the quantitative visualization of elasticity and viscosity based on molecular concentration. Brillouin images showed that the elasticity and viscosity of heterochromatin were higher than those of euchromatin and lower than those of the nucleolus. This result, together with Raman imaging, indicated that heterochromatin contains four times more nucleosomes, has a high lipid content, and is harder than euchromatin. The deviations in elasticity and viscosity are expected to be < 10%. One of the sources of the deviation is the changes in the peak position and bandwidth of the Brillouin spectra caused by the use of a high-numerical-aperture (NA) objective lens^50^, which arises from the fact that the Brillouin spectra depend on the angle of incident and scattered light. In our experimental setup with an objective lens (NA = 1.23), measurements with gelatin gel indicated that the shift in the peak position was < 0.2 GHz and bandwidth broadening was < 0.1 GHz at least when the bandwidth was ˃1 GHz (Supplementary Fig. 19). Thus, the use of a high-NA objective lens minimally impacted the obtained elasticity and viscosity images.

### Trichostatin A (TSA) treatment reduces the lipid concentration in heterochromatin and softens it

Next, we performed Raman and Brillouin imaging of live cells treated with TSA (Fig. 4). TSA is a potent histone deacetylase inhibitor that promotes the accumulation of highly acetylated histones, resulting in the relaxation of heterochromatin into euchromatin^51,52^. Lipid-rich regions corresponding to heterochromatin were observed in the Raman images of the nuclei of TSA-treated cells (Fig. 4a). DAPI fluorescence images revealed that the DNA concentration in heterochromatin was higher than that in the surrounding nucleoplasm (Supplementary Fig. 20). However, comparison of the Raman spectra with and without TSA treatment showed that treatment with TSA reduced the concentrations of DNA and lipids within heterochromatin, although the concentration of proteins remained almost unchanged (Fig. 4c, d). The DNA concentration decreased by approximately 30%, and the lipid concentration exhibited a more substantial decline, exceeding 60% in heterochromatin. The corresponding Brillouin image also captured TSA-induced alterations in heterochromatin (Fig. 4a). Adding TSA to the medium resulted in lower frequency shifts of the Brillouin bands accompanied by the bandwidth narrowing, indicating the softening of the nucleus (Fig. 4e-h). Treatment with TSA resulted in uniform spatial variation in the Brillouin image. In the absence of TSA treatment, heterochromatin was discernible in Brillouin images as regions exhibiting a broader band at a higher frequency than the surrounding nucleoplasm. Adding TSA reduced the differences in peak position and bandwidth between heterochromatin and the surrounding nucleoplasm, rendering them nearly indistinguishable in Brillouin images. TSA treatment reduced the peak position and bandwidth of the Brillouin band of heterochromatin, resulting in the softening of heterochromatin progressed to the same degree as the softening of euchromatin (Fig. 4g, h). This result is consistent with the function of TSA to induce the relaxation from heterochromatin to euchromatin^51,52^. As mentioned above, the Raman intensity analysis of TSA-treated cells revealed a 30% decrease in the DNA concentration in heterochromatin; however, the DNA concentration remained markedly higher than that in euchromatin. In contrast, the lipid content in heterochromatin decreased by 60%, suggesting that lipids play an essential role in controlling the physical properties of heterochromatin.

**Fig. 4.**
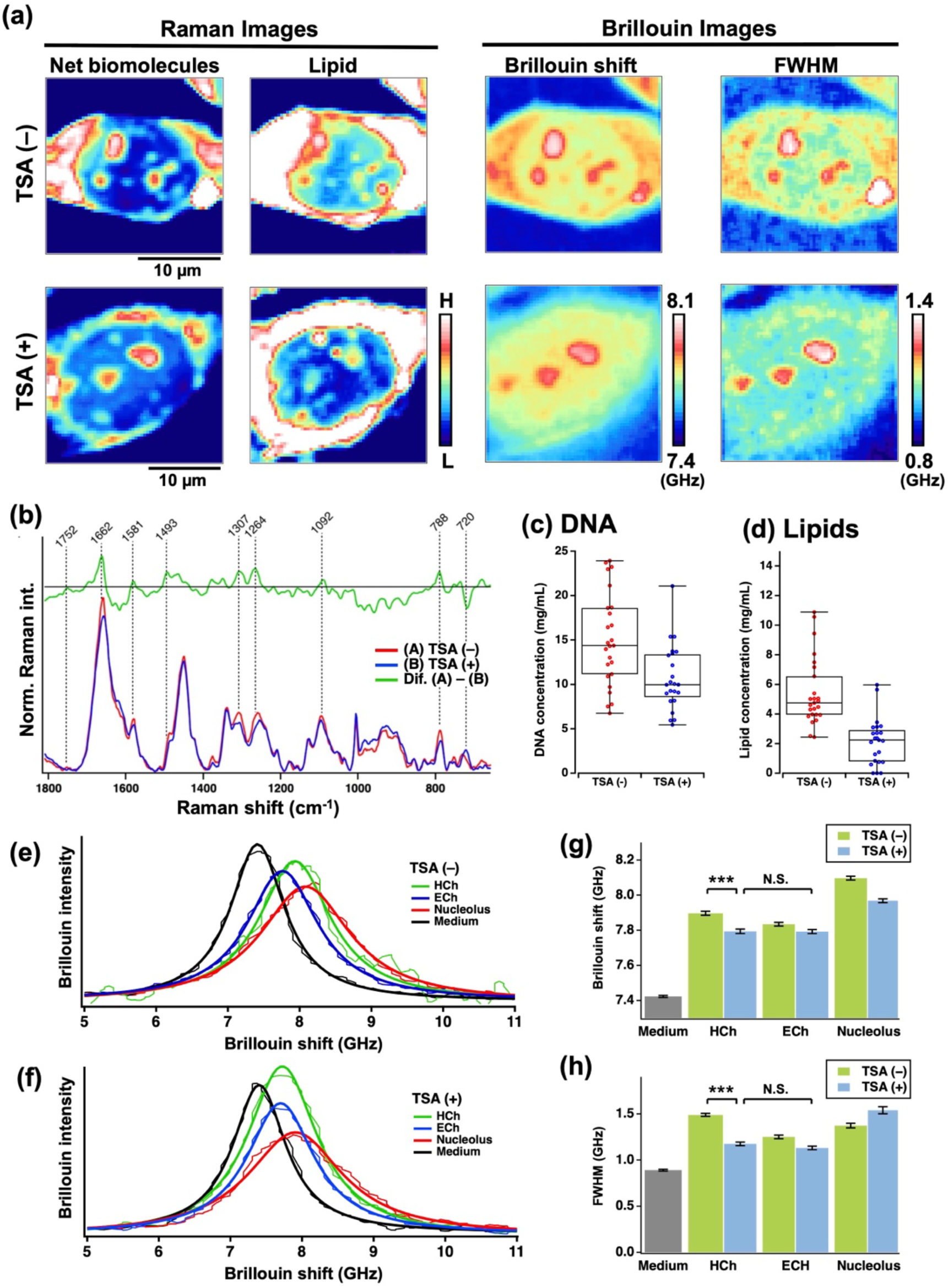
Chemical and physical changes in heterochromatin after TSA treatment. (a) Raman and Brillouin images of NIH3T3 cells with and without TSA treatment. Raman images were obtained by mapping the integrated intensity of the entire and low wavenumber regions of the C–H stretching band, visualizing the concentration distribution of the net biomolecules and lipids, respectively. Brillouin images were obtained by mapping the peak position and bandwidth at each pixel. (b-d) Raman spectral changes (b, (*n* = 6 for TSA (–), 4 for TSA (+))) and alternations in DNA (c) and lipid (d) concentrations in the chromocenter (*n* = 12). (e-h) Averaged (thin lines) and fitted (thick lines) Brillouin spectra before (e, *n* = 6) and after (f, *n* = 4) TSA treatment and changes in the peak position (g) and bandwidth (h) in heterochromatin (*n* = 10 for TSA (–), 13 for TSA (+)), nucleoplasm (*n* = 14 for TSA (–), 13 for TSA (+)), and nucleolus (*n* = 13 for TSA (–), 15 for TSA (+)). *n* represents the number of cells measured. *** : *p* < 0.001, N.S.: not significant.

### Chromosomes in mitosis do not contain lipids, and lipids are involved in the early G1 phase

We obtained a series of Raman and Brillouin images of NIH3T3 cells at different stages of mitosis, and captured the chemical and physical changes in chromosomes, which serve as the foundation for heterochromatin and euchromatin in interphase. (Fig. 5; representative Raman and Brillouin spectra and corresponding MCR spectra are shown in Supplementary Fig. 21 and 22, respectively). MCR images of DNA clearly show the characteristic behavior of chromosomes during mitosis: chromosomes begin to condense at prometaphase, align at the center of a cell at metaphase, split into sister chromatids, and are pulled toward opposite poles of the cell during the anaphase. In the subsequent telophase, chromosomes begin to decondense, and daughter nuclei are formed during cytokinesis. The corresponding Brillouin images exhibited almost identical features, indicating that condensed nucleosomes were stiffer than their surroundings. Changes in the chemical concentrations and Brillouin shifts in the chromosomes are summarized in Fig. 6. During anaphase, the DNA concentration in chromosomes was higher than that in other phases, as previously reported ^53,54,55^, with the Brillouin spectra exhibiting higher frequency shifts and broader bandwidths, thereby indicating that chromosomes in anaphase are harder than those in the other phases.

**Fig. 5.**
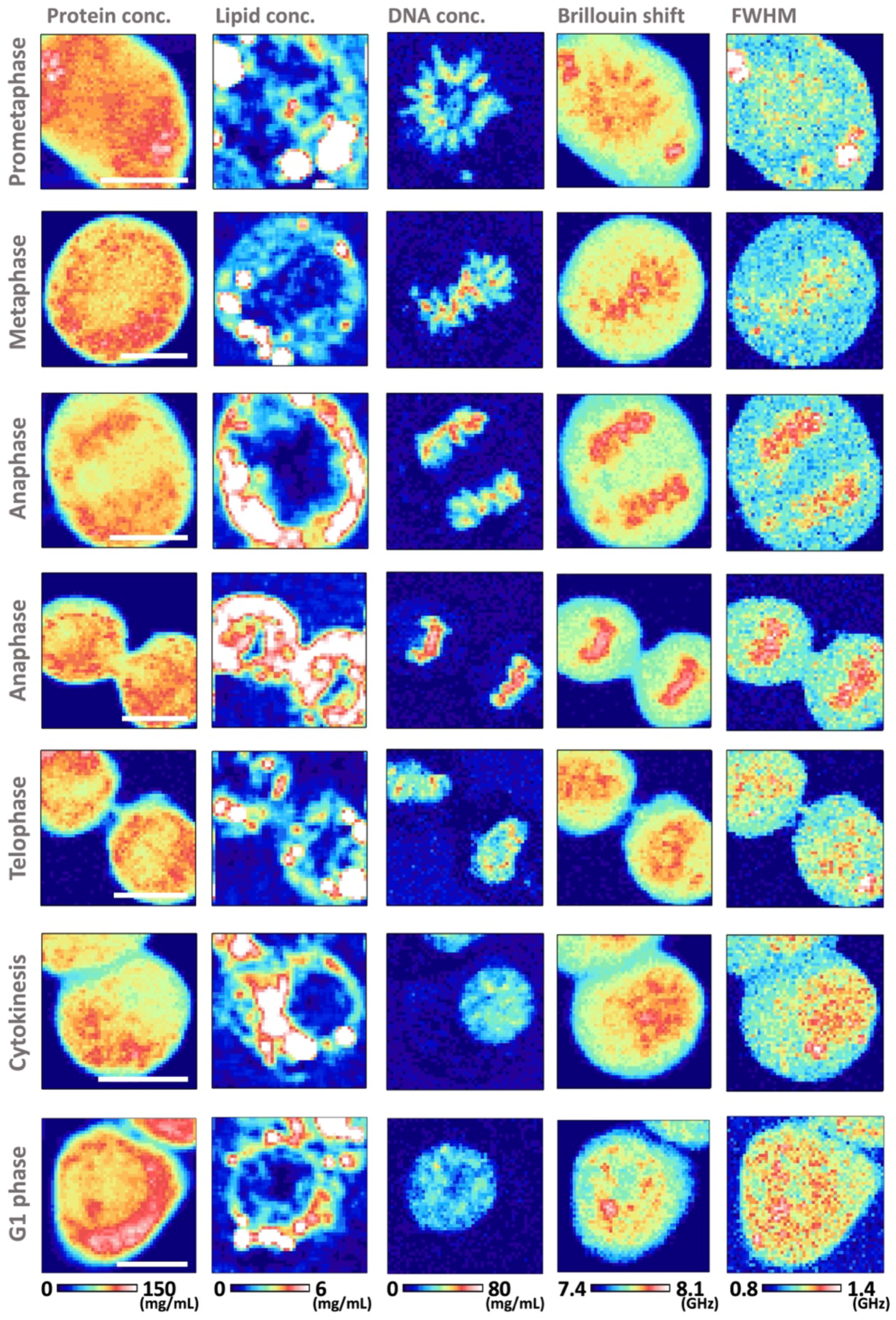
Raman and Brillouin images of NIH3T3 cells in mitosis. (Left to right) MCR images of proteins, lipids, DNA, and the corresponding Brillouin images. (top to down) NIH3T3 cells in prometaphase, metaphase, anaphase, anaphase, telophase, cytokinesis, and G1 phase. Brillouin images were obtained by mapping the peak position and bandwidth at each pixel. Scale bar, 10 μm.

**Fig. 6.**
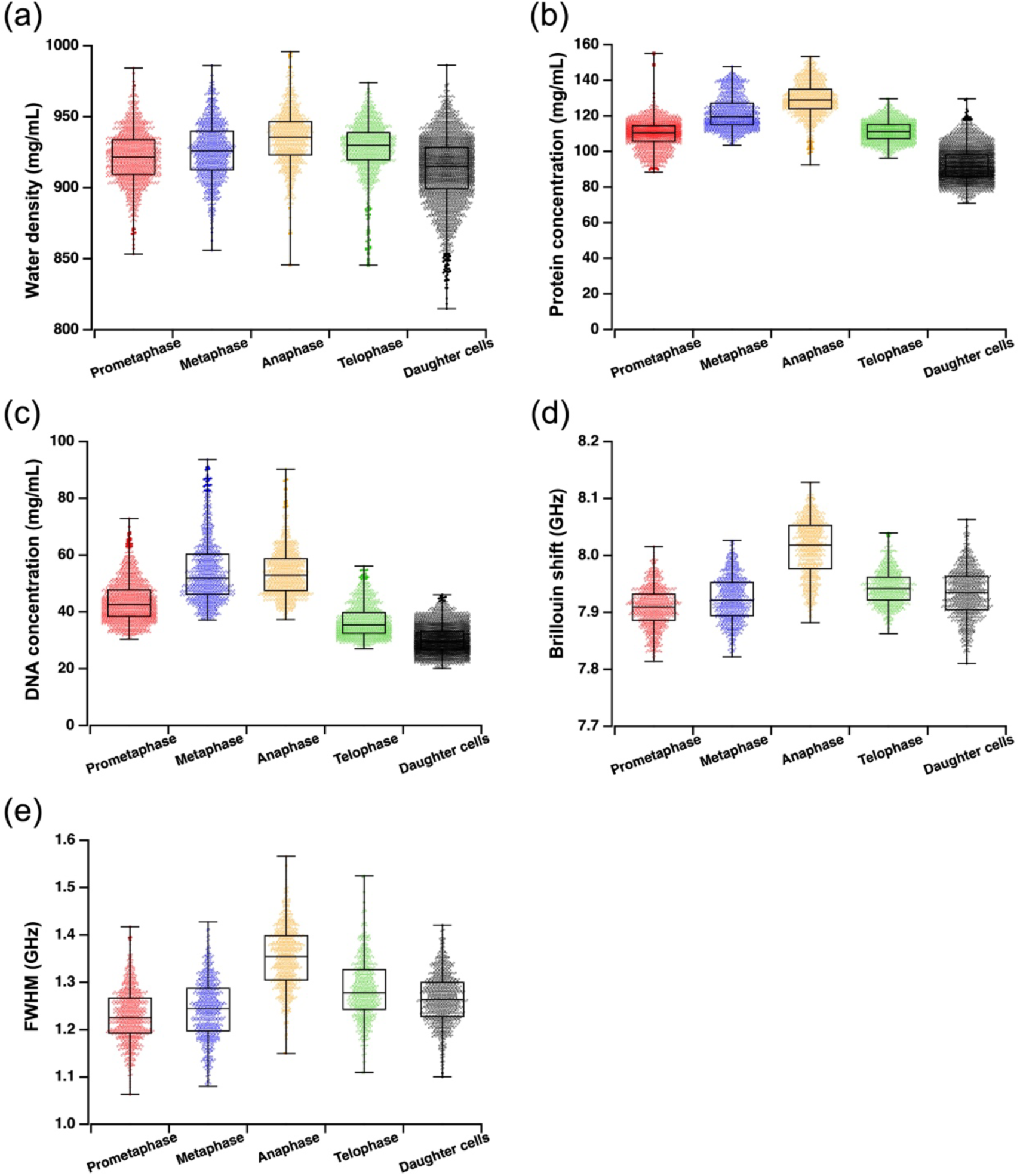
Chemical and physical changes in chromosomes of NIH3T3 cells during mitosis. (a-c) Concentration of water (a), proteins (b), and DNA (c) in chromosomes of NIH3T3 cells in each phase of mitosis, derived from Raman images. (d, e) Brillouin shift and the bandwidth of chromosomes in each phase. *n* = 5 for prometaphase, 5 for metaphase, 7 for anaphase, 5 for telophase, and 6 for daughter cells.

MCR images of lipids indicated the absence of lipids in chromosomes during mitosis. Even during telophase and cytokinesis, when chromosome decondensation and nucleus formation occur, MCR images of lipids showed no features in the nucleus, suggesting that no lipids are involved. In contrast, during the early G1 phase, some spots contained higher lipid concentrations in the nucleus (Fig. 7). Additionally, heterogeneous distributions of DNA and RNA with several DNA-enriched and RNA-enriched regions were observed (Fig. 7d, e). Comparing the lipid and nucleic acid images revealed that some DNA-enriched regions contained lipids. Analyses using the difference spectra indicated that DNA concentrations in DNA-only enriched regions were several times higher than those in the surrounding nucleoplasm. Furthermore, DNA/lipid-enriched regions contained slightly higher DNA concentrations than DNA-only enriched regions (Fig. 7g).

**Fig. 7.**
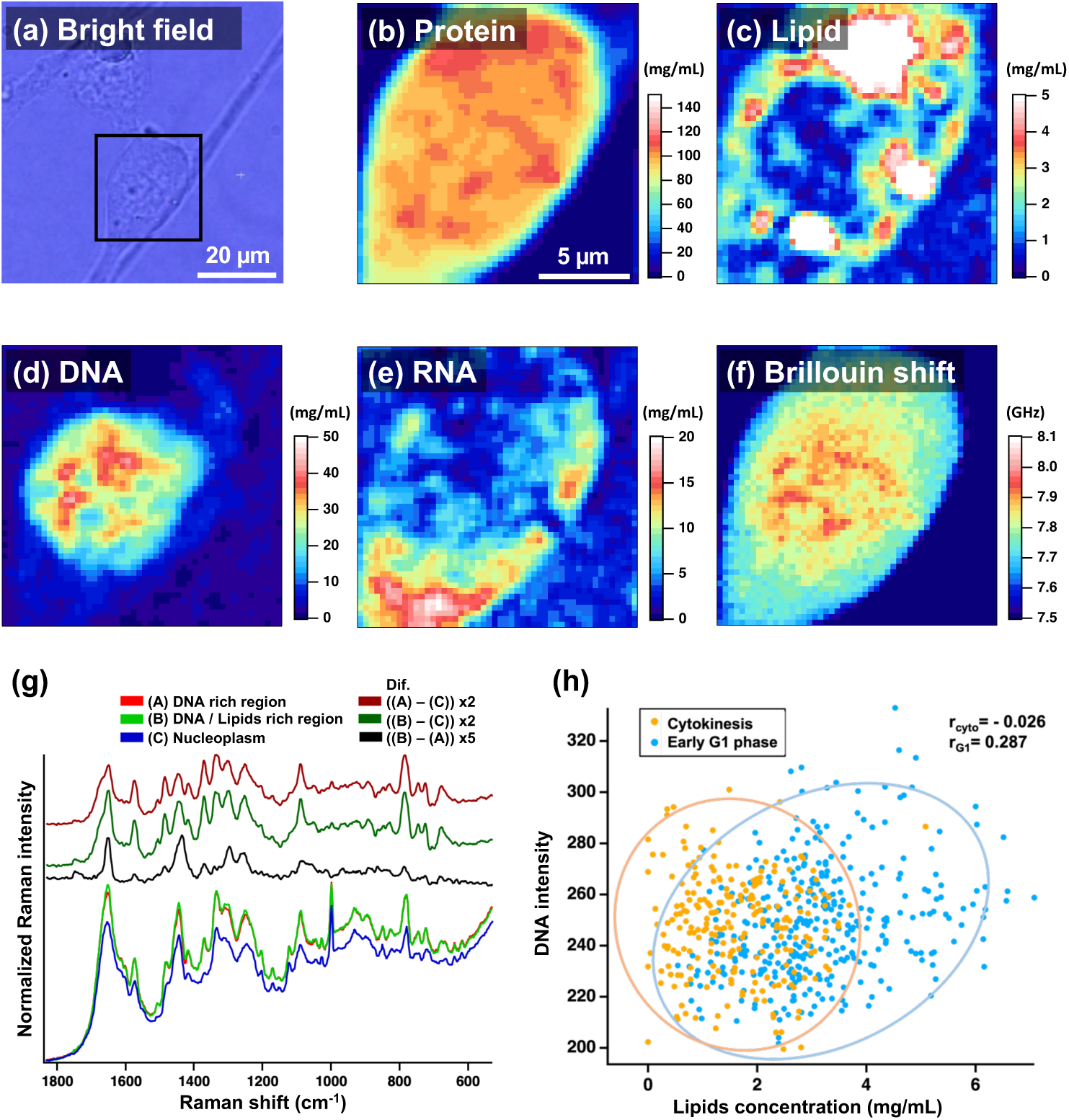
Lipid influx during heterochromatin formation in the early G1 phase. (a-f) Bright-field (a) and MCR images of proteins (b), lipids (c), DNA (d), RNA (e), and the corresponding Brillouin image of an NIH3T3 cell in the early G1 phase. Two daughter cells of similar size were attached to each other. (g) Raman spectra of DNA-rich and DNA/lipid-rich chromatin and the surrounding nucleoplasm region (*n* = 4 cells). The band in the pyrimidine band region (770-800 cm^-1^) of the black difference spectra ((B) - (A)) indicates a higher concentration in the DNA/lipid region than in the DNA-only region. (h) Correlation between the concentration of lipids and the Raman band intensity of DNA in the nucleus during cytokinesis (orange, *n* = 4 cells) and early G1 phase (blue, *n* = 4 cells).

Figure 7h shows the correlation between the Raman band intensities of lipids and DNA in the DNA-rich regions during cytokinesis and the early G1 phase. During the cytokinesis phase, there was only a small amount of lipids, with no correlation between lipids and DNA. Conversely, in the early G1 phase, a positive correlation was detected: the higher the lipid concentration, the higher the DNA concentration.

## Discussion

Herein, we developed a Raman and Brillouin microscopy system that offers simultaneous Raman and Brillouin imaging of a living cell. MCR-ALS analysis using a water Raman band as an internal standard enables the quantitative visualization of the concentration of each biomolecule and the total density distribution within a living cell. Combining this Raman approach with simultaneous Brillouin imaging makes it possible to quantitatively discuss the distributions of elasticity and viscosity within a cell based on the molecular concentrations. In this study, we used this method to analyze changes in heterochromatin under various conditions quantitatively: direct comparisons between heterochromatin and euchromatin, drug treatment, and cell cycle progression. We also quantitatively demonstrated that a certain amount of lipid molecules is deeply involved in heterochromatin formation.

MCR Raman analysis of NIH3T3 cells revealed that heterochromatin contained high concentrations of DNA and lipids. The DNA concentrations in heterochromatin and euchromatin were 15 and 4 mg/mL, respectively, indicating an approximate four-fold difference. Although several *in vitro* studies involving DNA extracts have suggested the presence of lipids in chromatin^56,57,58^, to the best of our knowledge, this is the first report on the direct observation of lipids in heterochromatin within a living cell. MCR images of cells in interphase clearly demonstrate that lipids are involved in both perinuclear and intranuclear heterochromatin. Furthermore, reduced lipid content in heterochromatin following TSA treatment, along with heterochromatin softening, indicated that lipids play an essential role in the condensation of heterochromatin and determine its chemical and physical properties. One possible hypothesis is that lipids incorporated into chromatin render the chromatin environment hydrophobic and function as molecular glue.

Raman imaging was performed on live cells without fixation. Although Raman imaging of fixed and DAPI-stained cells can visualize heterochromatin by mapping the intensity of the pyrimidine band, as well as the resonant Raman band of DAPI (Supplementary Fig. 23), lipid enrichment in heterochromatin was not observed in fixed cells (Supplementary Fig. 24). This could be due to lipid elution during paraformaldehyde fixation. Thus, a living cell needs to be observed to detect lipid molecules in heterochromatin.

Lipid molecules can enter protein coacervates, including liquid droplets, via hydrophobic interactions^59^. A similar mechanism is likely involved in the incorporation of lipid molecules into heterochromatin during its formation process, as liquid droplets formed through liquid-liquid phase separation by nucleosome and HP1α are regarded as an in-vitro model of heterochromatin^60,61,62^. The lipid influx renders the interior of the chromatin hydrophobic and decreases the dielectric constant, increasing electrostatic interactions between DNA and proteins and promoting heterochromatin condensation. Although no lipids were observed in chromosomes during mitosis, regions with high DNA concentrations were either lipid-rich or lipid-free during the early G1 phase. This result implies that when chromosomes decompose into heterochromatin and euchromatin in the newly formed nucleus after cell division, certain chromosomes incorporate lipids to maintain high DNA concentrations and become heterochromatin.

The next step was to identify the lipid molecules in heterochromatin. Lipid molecules present in the nucleus serve as signaling molecules^63,64^. Phosphoinositides are involved in DNA repair ^65^, ^66^; however, the fluorescence images of immunostained phosphatidylinositol bisphosphate (PIP2) and DAPI did not show co-localization, even after DNA damage due to X-ray irradiation, suggesting that lipids involved in heterochromatin are not PIP2 (Supplementary Fig. 25). The nucleus has no substantial membrane organelles, and the overall lipid concentration is normally low. Hence, where do nuclear lipids originate? One candidate is nuclear envelope invaginations ^67,68^, ^69^. Holographic microscopy revealed an average of five tubular organelles with a high refractive index in the nucleus of interphase NIH3T3 cells, which are presumed to be nuclear envelope invaginations (Supplementary Fig. 26). These tubular organelles can facilitate molecular exchange between the cytoplasm and nucleus. Contact between tubular organelles and chromocenters was observed, suggesting that lipids are supplied to heterochromatin via nuclear membrane invagination.

MCR analysis of Raman spectra within nuclear regions revealed that the main components of lipids in heterochromatin had phosphatidylcholine head groups (Supplementary Fig. 14). The MCR spectrum assigned to the lipids exhibited a spectral profile closely resembling that of DOPC^41,42^. This result is consistent with the above discussion that nuclear membrane invaginations serve as the source of lipids. The ratio between bands at 1660 and 1442 cm^-1^ showed that the average degree of lipid saturation in the nucleus was 1.4 times higher than that of DOPC, indicating that they consisted of a pair of fatty acids with one and two double bonds, such as 18:1 (oleic acid) and 18:2 (linoleic acid). The intensity ratio between the two bands at 719 cm^-1^ due to the choline group and 1442 cm^-1^ in the MCR spectrum was approximately half that in the spectrum of pure DOPC, implying that the proportion of lipids with a phosphatidylcholine head group was approximately 50%, and different types of lipids were also involved.

Although there were substantial differences in both DNA and lipid concentrations between heterochromatin and euchromatin, protein concentrations remained relatively constant, and there were no marked differences in the total concentration of biomolecules.

The difference in protein concentration between the heterochromatin and euchromatin was only 10%. The Brillouin spectra of heterochromatin and euchromatin display only a single peak at 7.9 and 7.8 GHz, which are comparable to 14% and 12% gelatin gels, respectively. This result indicated that both heterochromatin and euchromatin are relatively soft and liquid-like structures, and there are no rigid structures larger than the wavelength of phonons interacting with the excitation light (200 nm in this experiment)^6, 49^.

In conclusion, quantitative Raman and Brillouin microscopy offers label-free visualization of heterochromatin in living cells and reveals the involvement of lipids in heterochromatin formation. MCR analysis of Raman images of living NIH3T3 cells showed that the DNA concentration in heterochromatin was four times higher than in euchromatin, and heterochromatin comprised lipids with a phosphatidylcholine head group. Treatment with TSA reduced the amount of lipids, rather than DNA, in heterochromatin, rendering the heterochromatin softer. This result indicated that the amount of lipids is crucial for determining the physical properties of heterochromatin. Lipids were not observed in chromosomes during mitosis or the cytokinesis phase, indicating that heterochromatin acquires lipids during its formation as chromosomes are decondensed after nuclear membrane formation. Observations using a holographic microscope suggested that nuclear membrane invaginations are the lipid source. Lipid molecules play various roles in cells, including cell and organelle membrane formation, compartmentalization, and signal transduction. Our findings shed light on a new role for lipids in the nucleus: alternating the local environment to a more hydrophobic state and maintaining a high DNA concentration in heterochromatin, acting as a molecular glue.

## Methods

### Cell culture

NIH3T3 and HeLa cells were cultured in a glass-bottom dish (3960-035, 35 mm dish diameter, Matsunami) with 2 mL of culture medium (Dulbecco’s modified Eagle’s medium [DMEM] [D1145; Sigma] supplemented with 10% fetal bovine serum [10437-028; Gibco], 5×10^4^ U/L penicillin G, and 50 mg/L streptomycin sulfate [15070-063, Gibco]. The cells were incubated overnight at 37°C in a 5% CO2 humidified atmosphere. For the TSA experiment, after overnight incubation in a glass-bottom dish, the culture medium was replaced with DMEM containing 500 ng/mL trichostatin A, and the cells were incubated for an additional 24 h. Before Raman and Brillouin measurements, the medium was replaced with Hank’s balanced salt solution (HBSS).

### Raman Brillouin imaging

Raman and Brillouin images were acquired using a confocal Raman and Brillouin system (Nanofinder flex2, Tokyo Instruments Inc.), which integrated an inverted microscope (Eclipse Ti2, Nikon) with a Raman detection system (polychromator [MS3504i; SOL Inc.], EMCCD [DU970P-SVF, Andor]), and a Brillouin detection system (LightMachinery Inc.). A 532 nm excitation beam from a DPSS laser (Sprout Solo, Lighthouse Photonics) was focused onto the sample through a water immersion objective lens (Plan Apo IR 60XC/NA = 1.27, Nikon), which was used to collect both Raman and Brillouin scattering signals. The laser intensity at the entrance of the objective lens and exposure time for each pixel were 40 mW and 0.2 s, respectively. The sample stage was scanned at intervals of 300 nm using a piezo stage (NanoControl) to acquire Raman and Brillouin images. Interphase cells were measured at 37℃ using a stage-top incubator (STXF-TIZBX-SET, Tokai hit), while mitotic cells were measured at room temperature. All Raman and Brillouin images were analyzed using the IGOR Pro program package (WaveMetrics). Singular value decomposition (SVD) was applied to each Raman image to reduce noise. Raw Raman spectral images were used for the MCR analysis to ensure that small bands were not overlooked. Prior to each MCR analysis, the region of a strong and sharp phenylalanine band at around 1,000 cm^-1^ was omitted to prevent interference with the MCR component spectra. Each Brillouin spectrum was fitted with a Lorentzian function, and the peak position and bandwidth were obtained as averages of the Stokes and anti-Stokes bands.

### DAPI fluorescence imaging

After Raman and Brillouin measurements, cells were fixed with 4% paraformaldehyde on the microscope stage and stained with DAPI containing DMEM at 37℃ for 30 min. The medium was then replaced with HBSS, and fluorescence imaging was performed using the same objective lens and a CMOS camera (Wex120, Wraymer). Images obtained were processed using ImageJ software (National Institutes of Health, Bethesda, MD).

### Immunofluorescence imaging of cells with ionizing radiation

Retinal Pigment Epithelium (RPE) cells were cultured in a glass-bottom dish (35 mm dish diameter, 3960-035; Matsunami) with 2 mL of culture medium (DMEM [Sigma] supplemented with 10% fetal bovine serum [Gibco], 5×10^4^ U/L penicillin G, and 50 mg/L streptomycin sulfate [Gibco]). After 48 h of incubation at 37°C under a 5% CO2 humidified atmosphere, the cells were irradiated with 1 Gy of X-ray radiation from an X-ray light source (MX-160Labo, MediXtec) at 160 kV and 3 mA for 54 s. After an additional 30 min of incubation, cells were fixed in phosphate-buffered saline containing 4% paraformaldehyde and 2% sucrose at room temperature for 10 min. Subsequently, cells were immunostained for PIP2 (Anti-PIP2 antibody [2C11]; ab11039, Abcam), along with Mouse 488 (Anti-mouse IgG (H+L), F(ab’)2 Fragment [Alexa Fluor 488 Conjugate]; 4408S, Cell Signaling Technology) and 53BP1 (Anti-53BP1 antibody [Rabbit polyclonal]; A300-272A, Bethyl); and Rabbit 594 (Anti-rabbit IgG (H+L), F(ab’)2 Fragment [Alexa Fluor 594 Conjugate]; 8889S, Cell Signaling Technology). After mounting cells with ProLong Gold (ProLong Gold Antifade Reagent with DAPI) (8961S; Cell Signaling Technology), fluorescence images were obtained using a cooled CMOS camera (WRAYCAM-CIX830, Wraymer).

### Refractive index measurements

Cells were cultured in a glass-bottom dish (Tomodish, Tomocube) and incubated overnight at 37°C in a 5% CO2 humidified atmosphere. After replacing the culture medium with HBSS, 3D refractive-index imaging was performed using a holographic microscope (HT-2H; TomoCube).

### Raman measurements of solution

A homogeneous solution of yeast RNA (AM7118; Invitrogen), DNA (D1626; Sigma), lysozyme (127-06724; Wako), pepsin (20569-02; Nacalai), bovine serum albumin (BSA) (08587-26; Nacalai), and oleic acid (159-00246; Wako) was used as a standard solution for the calibration lines. While DNA and RNA were dissolved or diluted in ultrapure water, proteins were dissolved in a buffer solution at pH 7.4. Oleic acid was mixed with n-decan (10613-25; Nacalai) in various concentrations. The concentrations of proteins and DNA/RNA were determined by measuring the absorption at 280 and 260 nm, respectively. The solution was placed on a glass-bottomed dish, and Raman spectra of the solution were measured using a Raman/Brillouin microscope. For the oleic acid/decan mixture, Raman images of oil droplets in a glass-bottom dish filled with water were obtained.

### Statistics and reproducibility

Statistical significance was determined by one-sided unpaired Student’s t-test and one-sided binomial test with Excel software (Microsoft, Redmond, WA). Differences were considered statistically significant at *p* < 0.05. All representative Raman/Brillouin images were reproduced the number of times indicated in the corresponding plot analyses and/or average spectra. No data were excluded from the analyses and no statistical method was used to predetermine the sample size.

## Acknowledgements

This work was supported by JSPS KAKENHI Grant Numbers JP24KJ04080 (MM), JP17H05869 (TN), JP21H05261 (TN), JP22H02594 (TN), JP19H02666 (SK), JP20H04689 (SK), and JP24K02161 (SK) from the Ministry of Education, Culture, Sports, Science, and Technology in Japan, and JST, PRESTO Grant Number JPMJPR20E5 (SK), CREST Grant Number JPMJCR2024 (TN). SK also acknowledges the research grants from Kurita Water and Environment Foundation (KWEF) (19E030), Uehara Memorial Foundation, and Nakatani Foundation in Japan.

## Author contribution

S.K., K.F., A.S., and T.N. conceived the idea. M.M., S.K., K.H., and T.N. designed the experiments. M.M. and S.K. performed Raman/Brillouin imaging and analyzed the data. M.M., S.K., and R.S. wrote MCR-ALS codes used in this study. Y.O. and M.H. assisted sample preparations. M.M., M.I., and A.S. performed immunofluorescence imaging of cells with ionizing radiation. M.M., M.W., and M.O. performed Refractive index measurements. All the authors checked the data and discussed the results. M.M., S.K., and T.N. wrote the manuscript. All the authors edited the manuscript. All the authors read and approved the final manuscript.

