## Supplementary Information for "Lipids Are Involved in Heterochromatin Condensation: A Quantitative Raman and Brillouin Microscopy Study"

**Corresponding authors**

This document contains the following information.

Supplementary Figures:

1. Chromocenters in an NIH3T3 cell.
2. Raman spectra of DNA and RNA.
3. Raman images of a living NIH3T3 cell.
4. MCR spectra of an NIH3T3 cell. .
5. Comparison between MCR spectra and reference spectra.
6. Reconstructed Raman spectra of different cellular compartments.
7. Raman spectra of proteins.
8. Raman spectra of oleic acid.
9. Raman spectra of DNA and RNA.
10. Calibration lines for the concentrations of biomolecules.
11. Co-localization of DNA and lipids in heterochromatin of an NIH3T3 cell.
12. MCR images of a HeLa cell.
13. Co-localization of DNA and lipids in heterochromatin of a HeLa cell.
14. MCR spectra for nucleus regions exclusively.
15. Phosphatidylcholine is the main component of lipids in the nucleus.
16. Raman spectra provide information on lipid unsaturation levels.
17. Refractive index of nuclear compartments.
18. Refractive index map of the NIH3T3 cell based on Raman imaging.
19. Effects of high-numerical-aperture (NA) objective lens on Brillouin spectra.
20. Chromocenters in trichostatin A (TSA)-treated cells.
21. Raman and Brillouin spectra of NIH3T3 cells in mitosis.
22. MCR spectra of mitotic and interphase NIH3T3 cells.
23. Raman images of a DAPI-stained NIH3T3 cell.
24. Raman spectra of DAPI-stained NIH3T3 cells.
25. Immunofluorescence images show that $PIP_2$ does not localize in the chromocenter.
26. Refractive index images of nuclear membrane invagination tubes.

Supplementary Table:

1. Assignments of Raman bands

Supplementary References

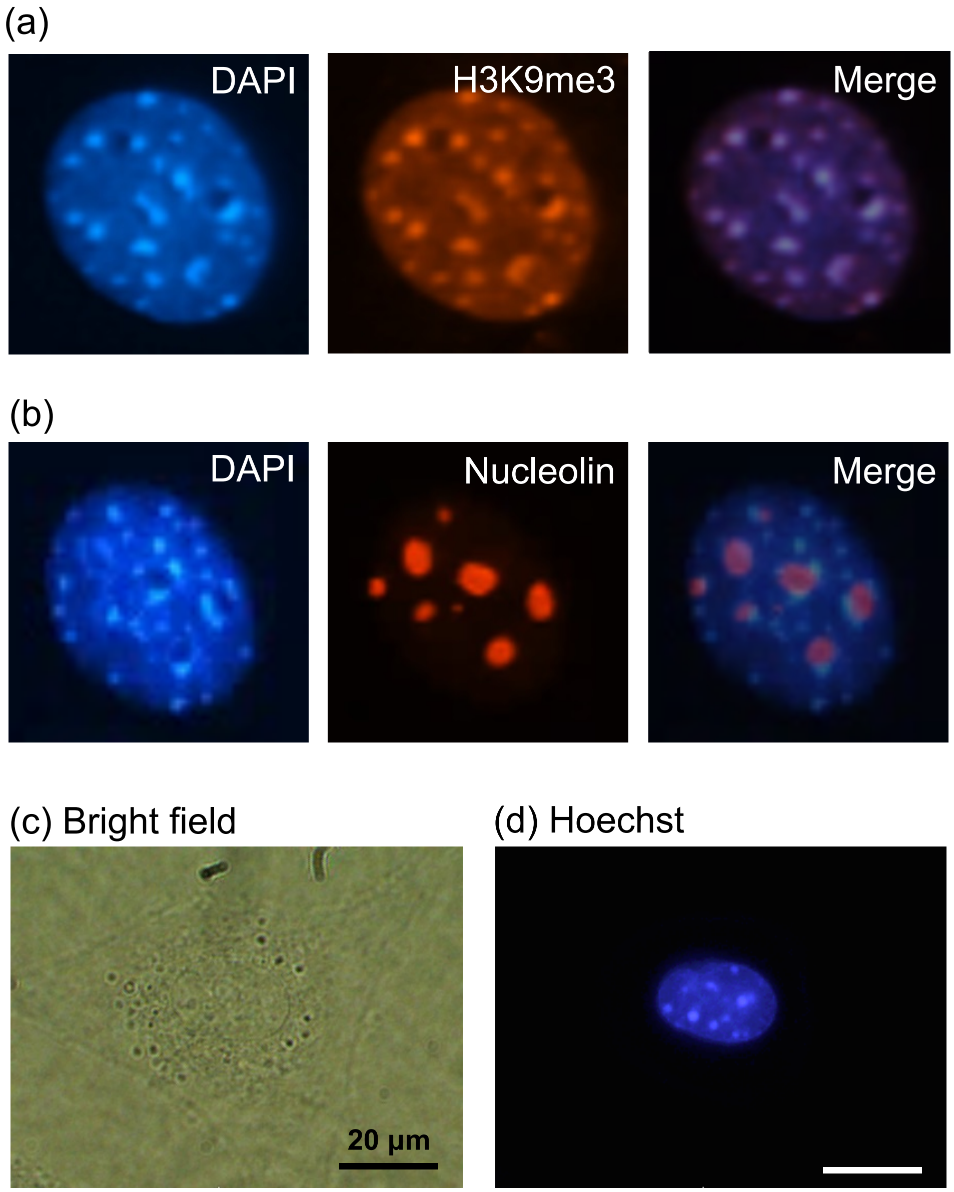


**Supplementary Fig. 1. Chromocenters in an NIH3T3 cell.** (a, b) Fluorescence images of a fixed NIH3T3 cell stained with DAPI and H3K9me3 (a) or DAPI and nucleolin antibody (b) and their merged images. H3K9me3 and nucleolin are markers for heterochromatin and nucleolus, respectively. Merged images demonstrate that DAPI-stained chromocenters correspond to heterochromatin. (c, d) Bright-field (c) and the corresponding Hoechst fluorescence (d) images of a living NIH3T3 cell. Panel (d) shows that chromocenters in living cells are visualized by Hoechst staining.

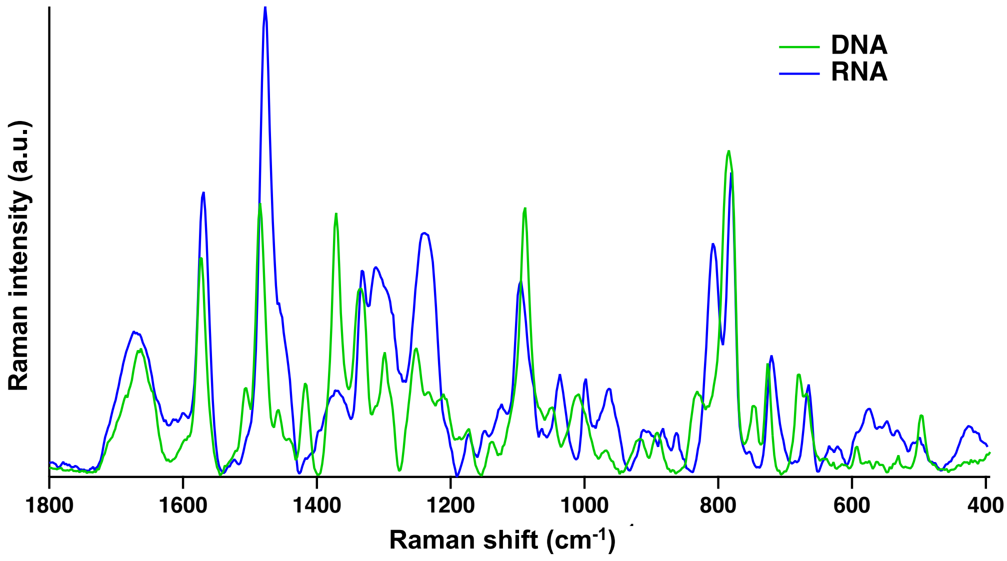


**Supplementary Fig. 2. Raman spectra of DNA and RNA.** Raman spectra of DNA (green) and RNA (blue). DNA was measured in an aqueous solution, while RNA was encapsulated in aqueous droplets formed in a polyethylene glycol (PEG) solution and measured. An aqueous solution of RNA was mixed with PEG powder, resulting in the formation of aqueous droplets containing a high concentration of RNA in 50% PEG solution[1].

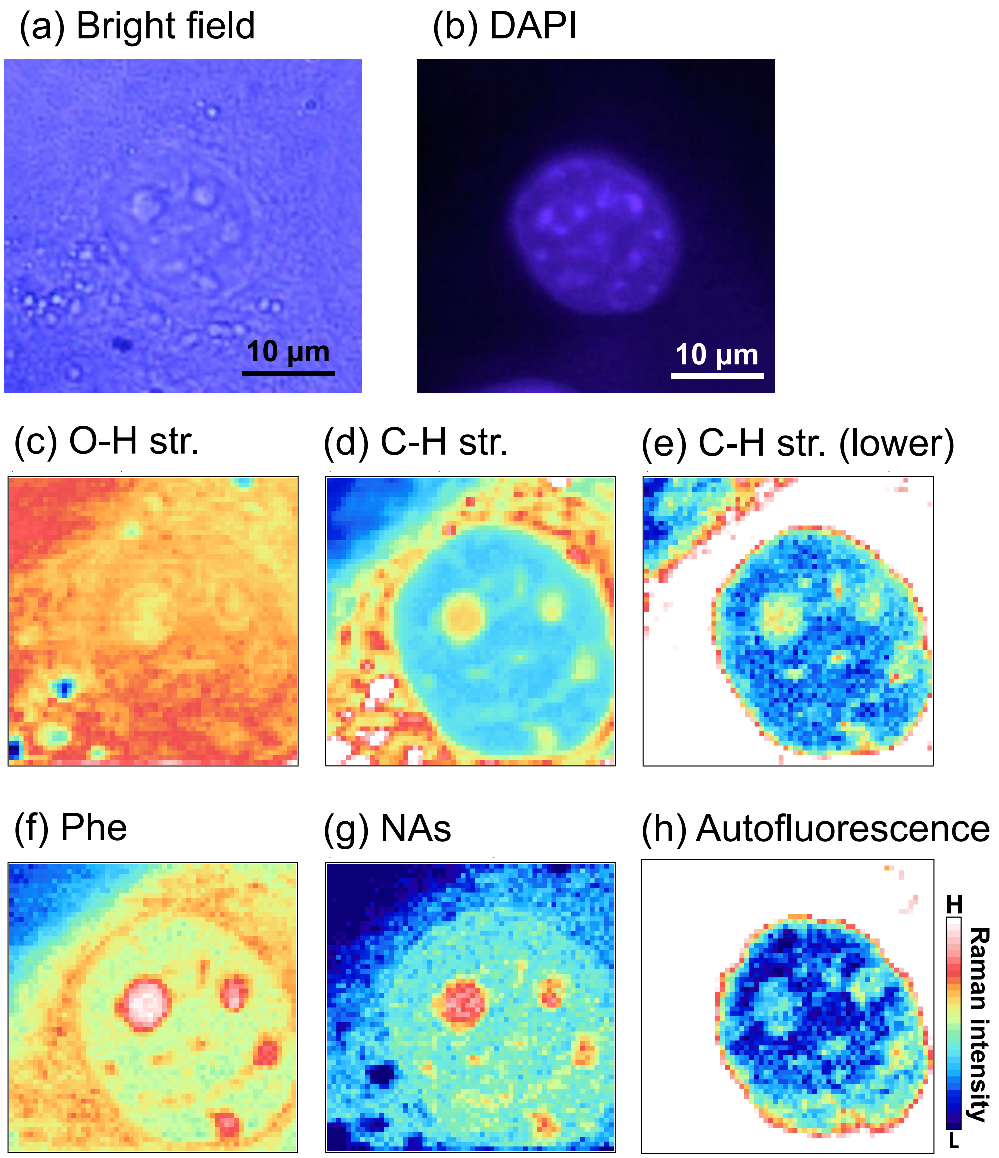


**Supplementary Fig. 3. Raman images of a living NIH3T3 cell.** (a-h) Bright-field (a), DAPI fluorescence (b), and conventional Raman (c-g) and autofluorescence (h) images of an NIH3T3 cell. The fluorescence image was obtained after obtaining Raman images, followed by cell fixation and staining with DAPI. The corresponding MCR images are shown in Fig. 2 (in the main text). Raman images were obtained by mapping the intensity of the O–H stretching band (c, 3500–3650 $cm^{-1}$), the entire region (d, 2800–3030 $cm^{-1}$) and the lower-wavenumber region (e, 2850–2890 $cm^{-1}$) of the C–H stretching band, phenylalanine band (f, 995–1010 $cm^{-1}$), and the pyrimidine band (g, 770–800 $cm^{-1}$). The autofluorescence image was obtained by mapping the intensity of the silence region of the Raman spectra (h, 2000–2400 $cm^{-1}$).

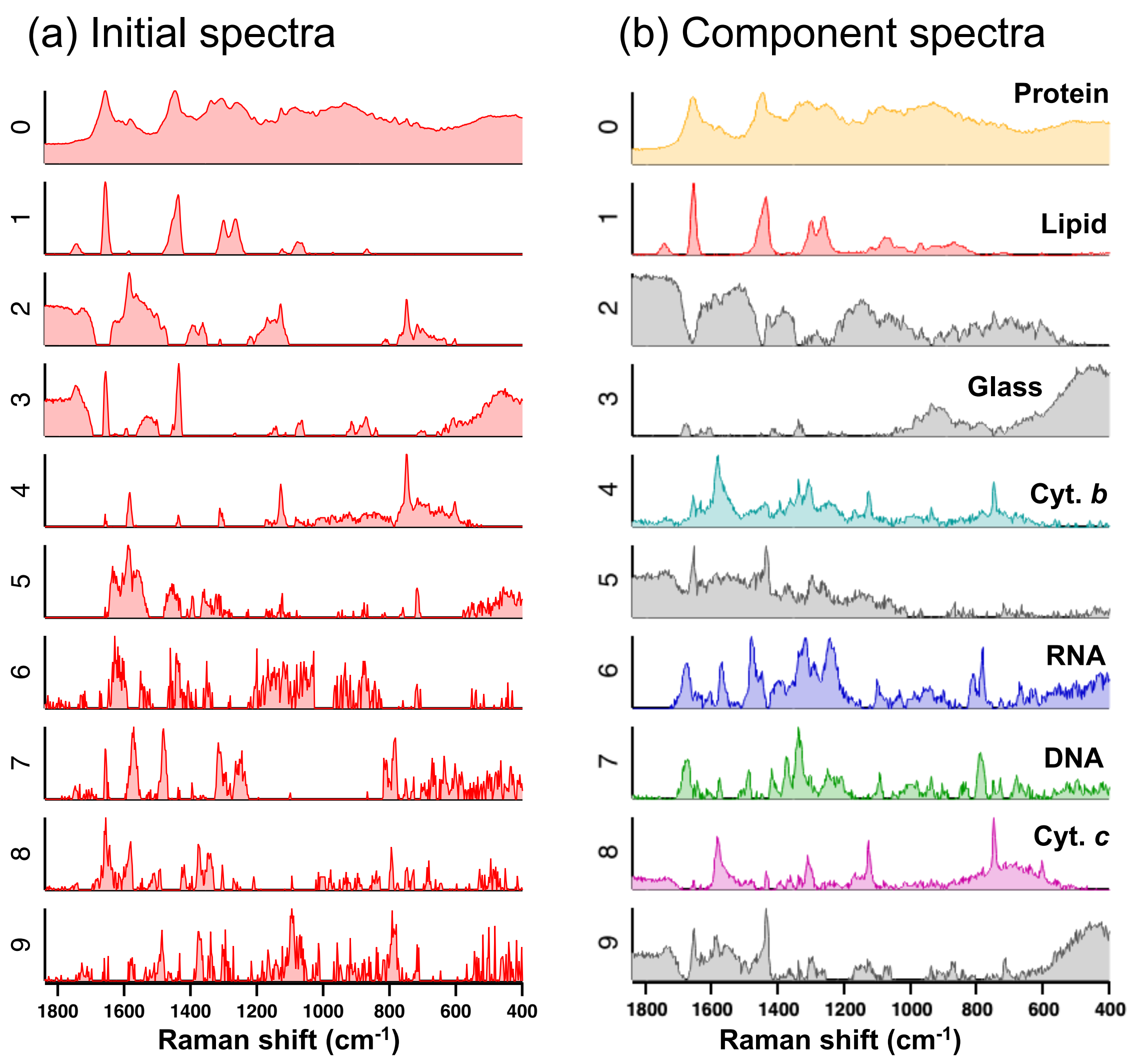


**Supplementary Fig. 4. MCR spectra of an NIH3T3 cell.** (a) Initial spectra for MCR analysis obtained by a singular value decomposition (SVD)-NMF (non-negative matrix factorization) method. (b) Component spectra obtained by the MCR analysis. The corresponding MCR and original Raman images are shown in Fig. 2 (in the main text) and Supplementary Fig. 3, respectively. Each component assignment, based on comparison with reference Raman spectra, is shown in the upper right corner of each panel.

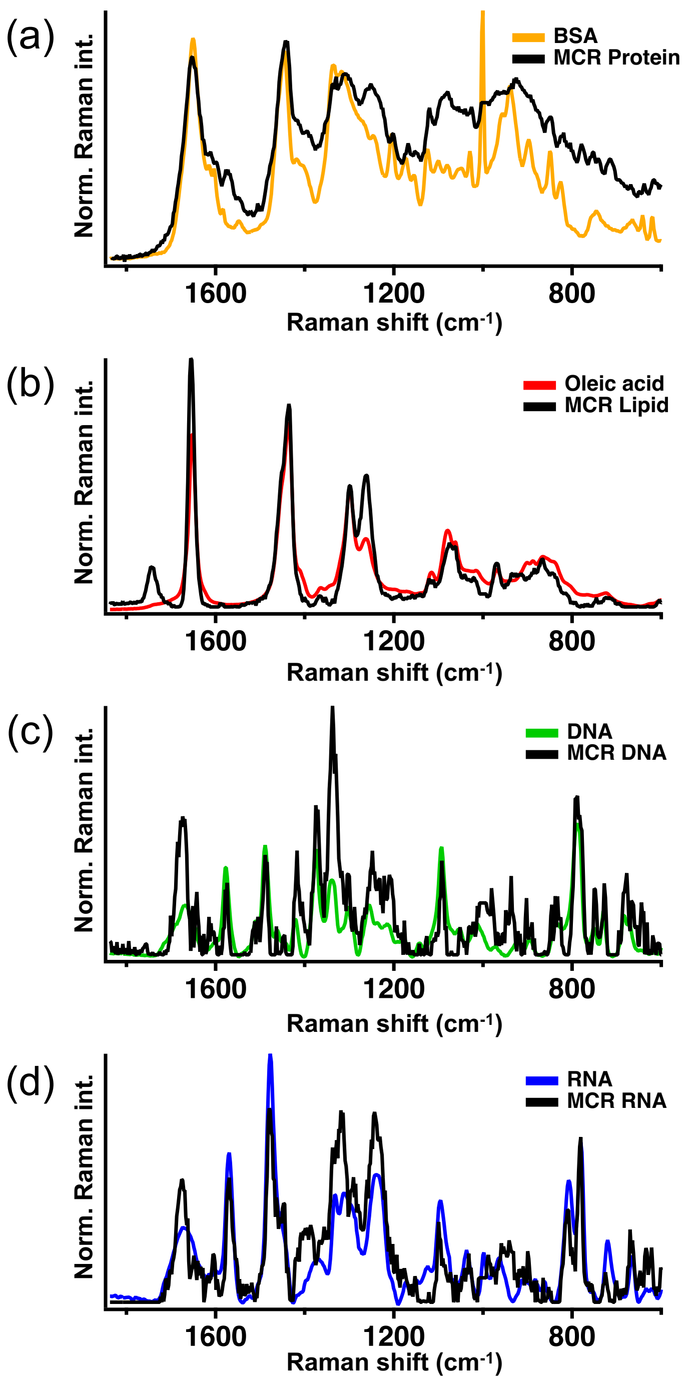


**Supplementary Fig. 5. Comparison between MCR spectra and reference spectra.** (a-d) Comparison of the MCR and Raman spectra of bovine serum albumin (BSA) (a), oleic acid (b), DNA (c), and yeast RNA (d).

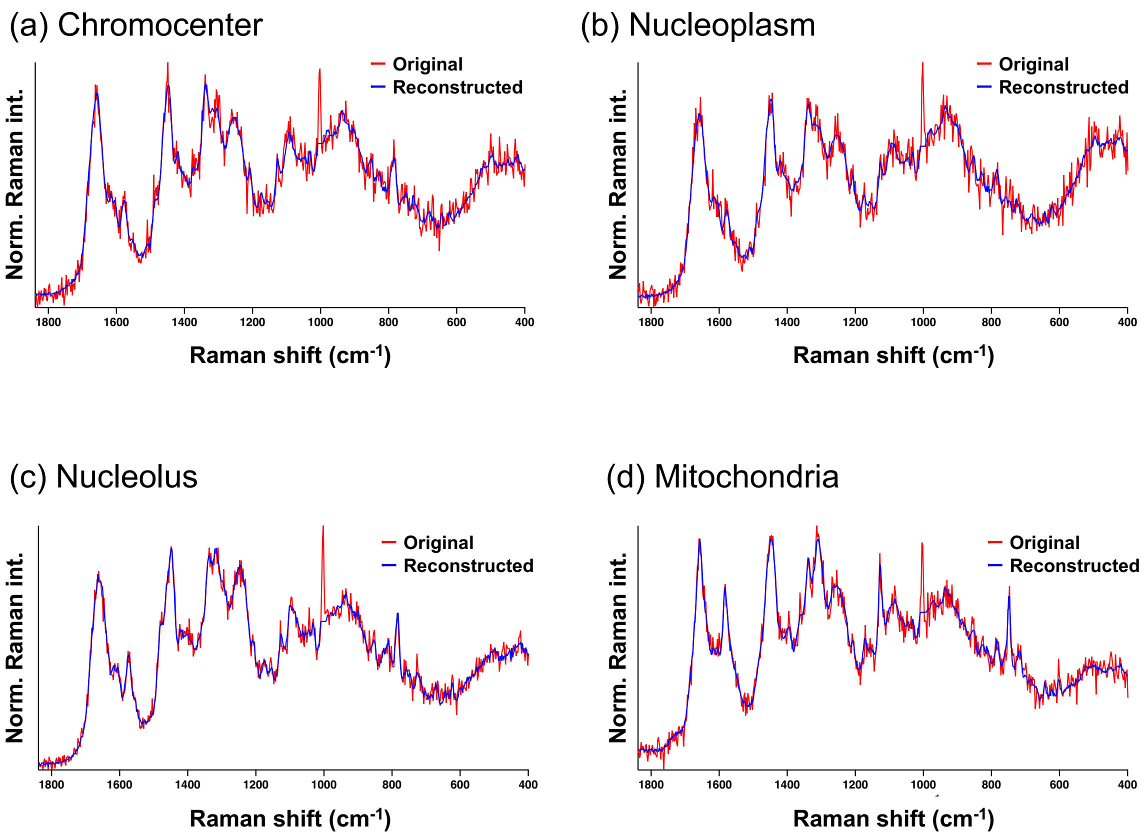


**Supplementary Fig. 6. Reconstructed Raman spectra of different cellular compartments.** (a-d) Original Raman and reconstructed spectra of chromocenter (a), nucleoplasm (b), nucleolus (c), and mitochondria (d). After MCR analysis, reconstructions were performed using the MCR spectra of 6 main components (proteins, RNA, DNA, lipids, and cytochrome *c* and *b*) and glass substrate and their distribution image.

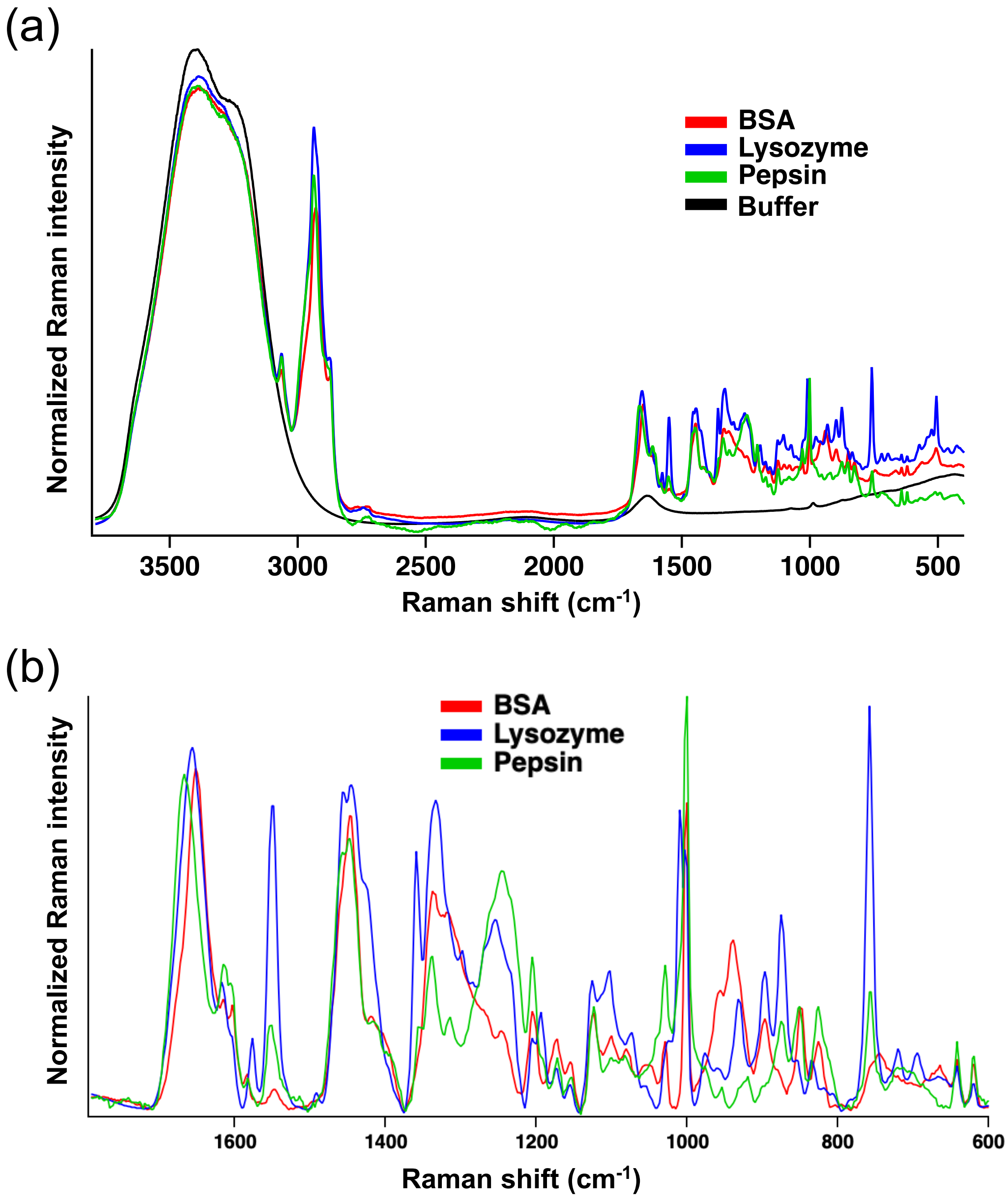


**Supplementary Fig. 7. Raman spectra of proteins.** (a, b) Raman spectra of bovine serum albumin (BSA), lysozyme, and pepsin in buffer solutions of the entire (a) and fingerprint (b) regions, along with the Raman spectrum of the buffer solution alone. Panel b shows spectra obtained by subtracting the Raman spectrum of the buffer solution.

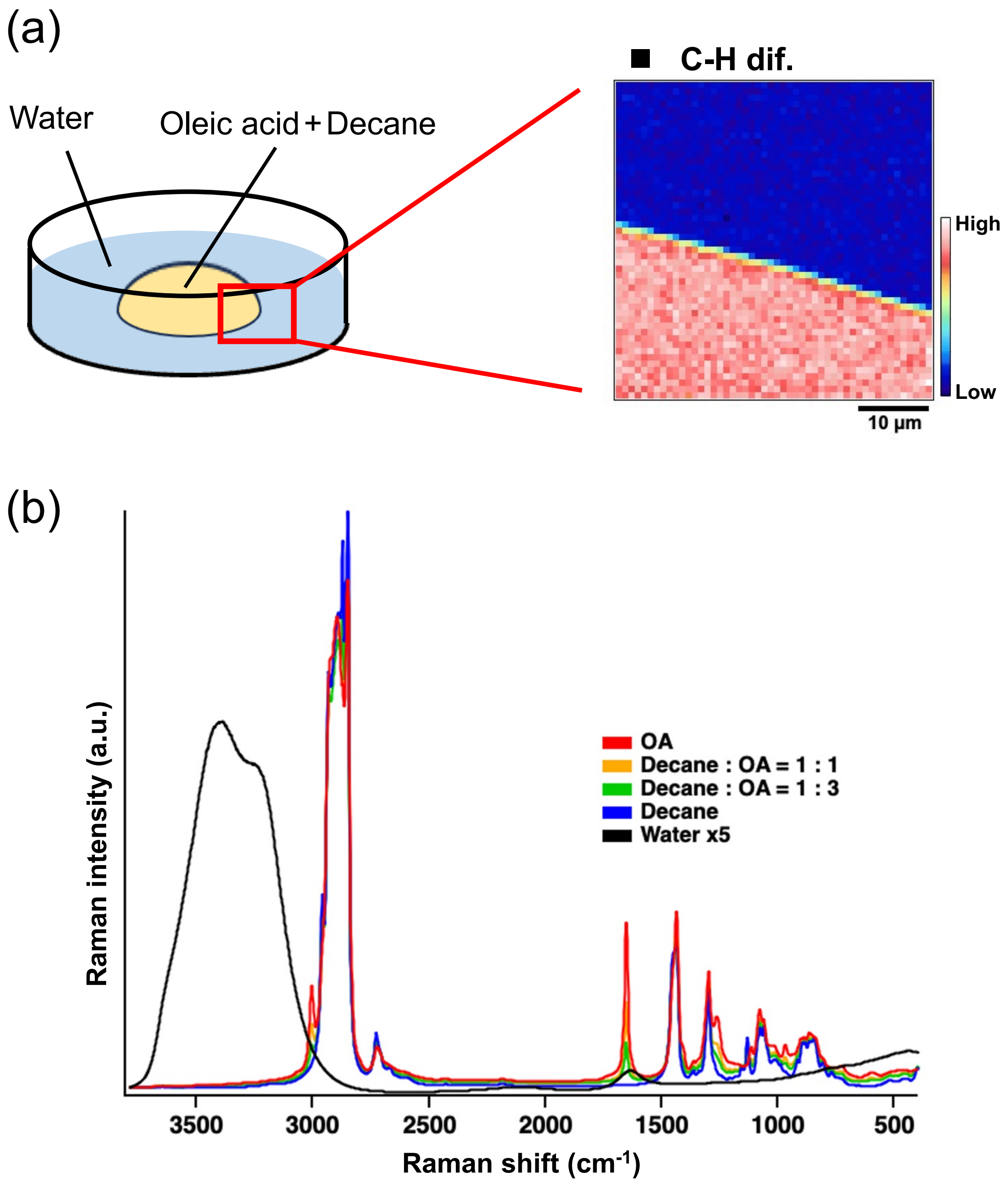


**Supplementary Fig. 8. Raman spectra of oleic acid.** (a) Schematic illustration of Raman imaging measurements of oleic acid (OA)/decane droplets in water. (b) Raman spectra of OA/decane droplets with varying OA concentrations, along with the Raman spectrum of the surrounding water.

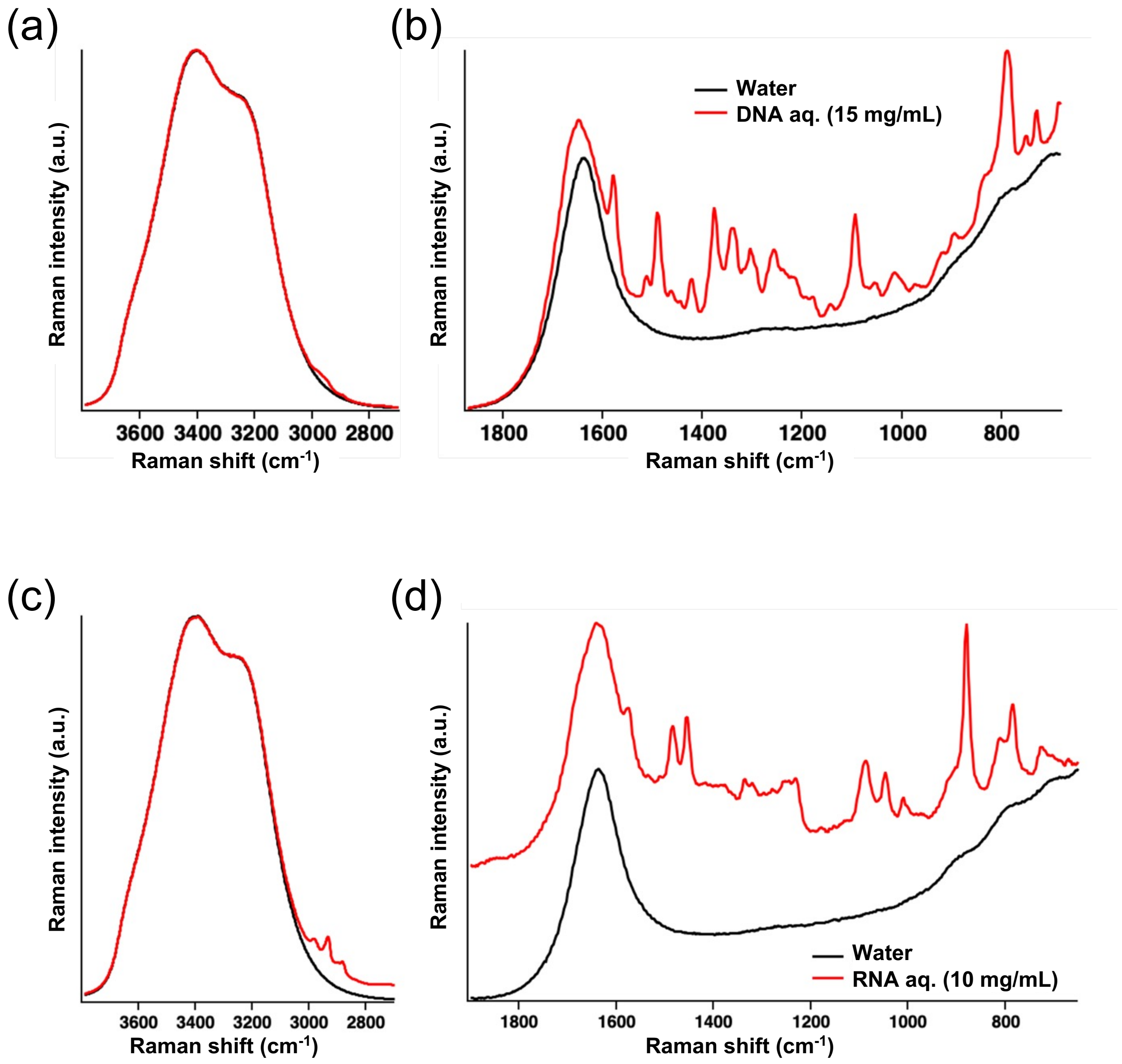


**Supplementary Fig. 9. Raman spectra of DNA and RNA.** (a-d) Raman spectra of 15 mg/mL DNA (a, b) and 10 mg/mL RNA solution (c, d) in the high-wavenumber (a, c) and fingerprint region (b, d), along with a Raman spectrum of pure water.

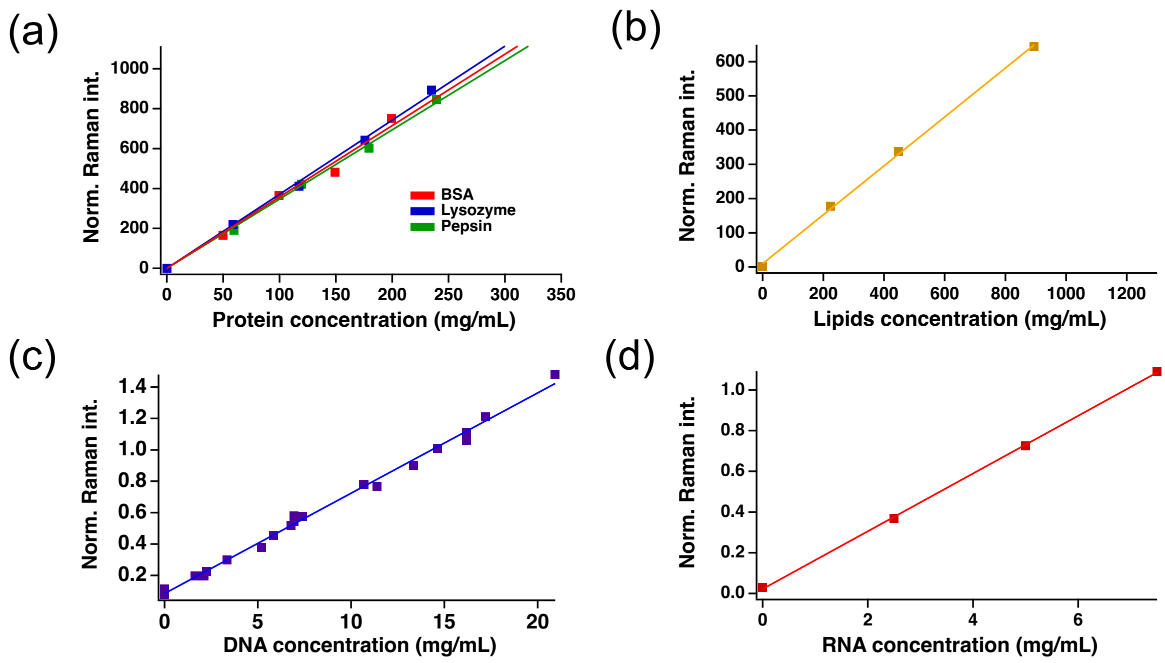


**Supplementary Fig. 10. Calibration lines for concentrations of biomolecules.** (a-d) Calibration lines for proteins (a), lipids (b), DNA (c), and RNA (d) obtained by plotting the Raman intensity of biomolecules relative to the intensity of the water O–H band (~ 3400 $cm^{-1}$) against their concentrations, using Raman bands at 1650 $cm^{-1}$ (amide I) for proteins, 1435 $cm^{-1}$ ($CH_2$ scissors) for lipids, and 790 $cm^{-1}$ (pyrimidine ring) for DNA/RNA.

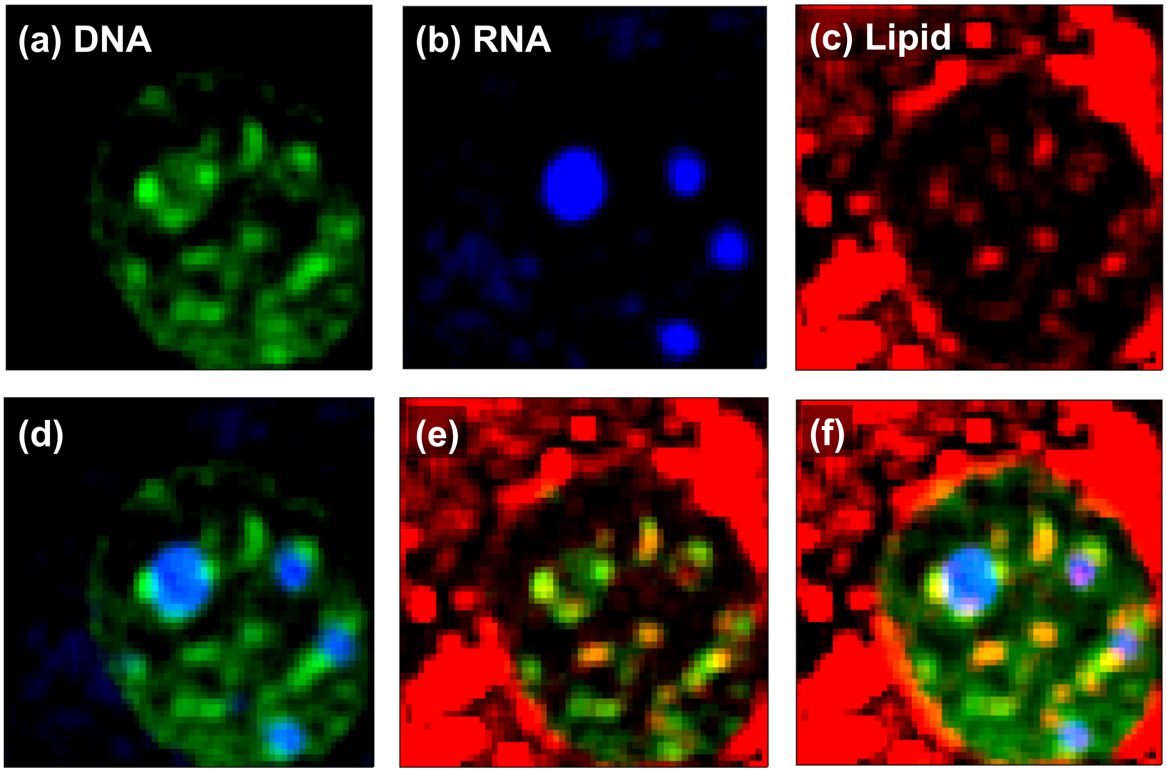


**Supplementary Fig. 11. Co-localization of DNA and lipids in heterochromatin of an NIH3T3 cell.** (a-f) MCR images of DNA (a, green), RNA (b, blue), lipids (c, red), and their merged images (d-f). Original MCR images are shown in Fig. 2 (in the main text).

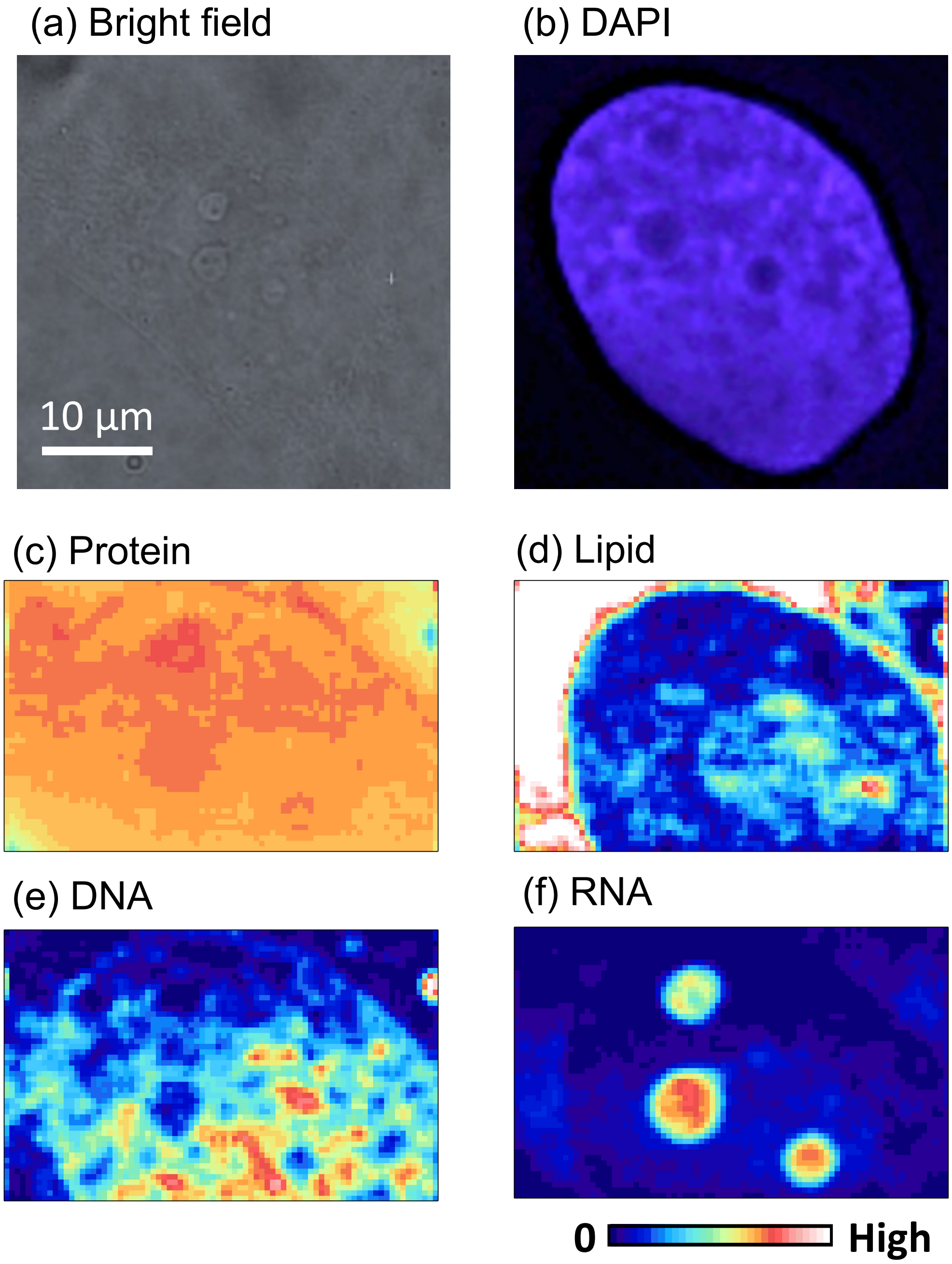


**Supplementary Fig. 12. MCR images of a HeLa cell.** (a-d) Bright-field (a), DAPI fluorescence (b), and MCR Raman (c-f) images of a living HeLa cell. Each MCR image corresponds to the distribution of proteins (c), lipids (d), DNA (e), and RNA (f), respectively.

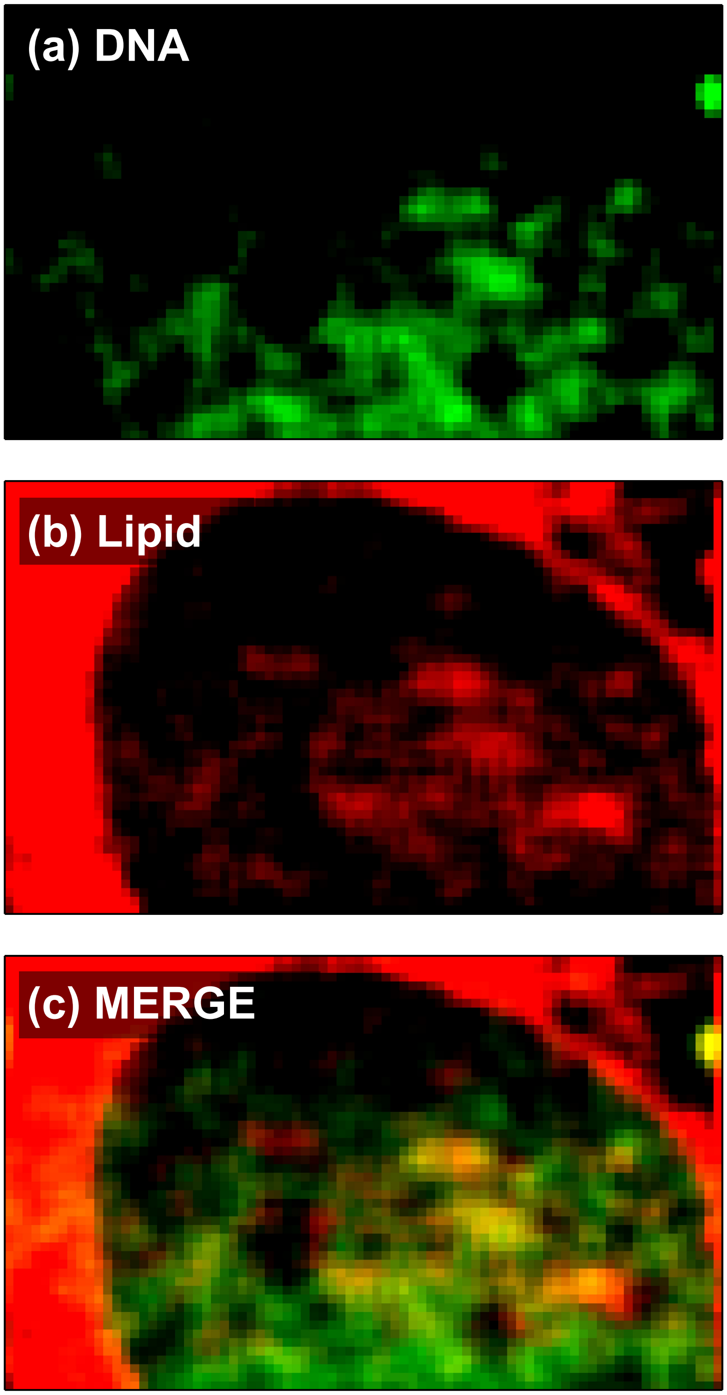


**Supplementary Fig. 13. Co-localization of DNA and lipids in heterochromatin of a HeLa cell.** (a-c) MCR images of DNA (a, green) and lipids (b, red) and their merged images (c). Original MCR images are shown in Supplementary Fig. 12.

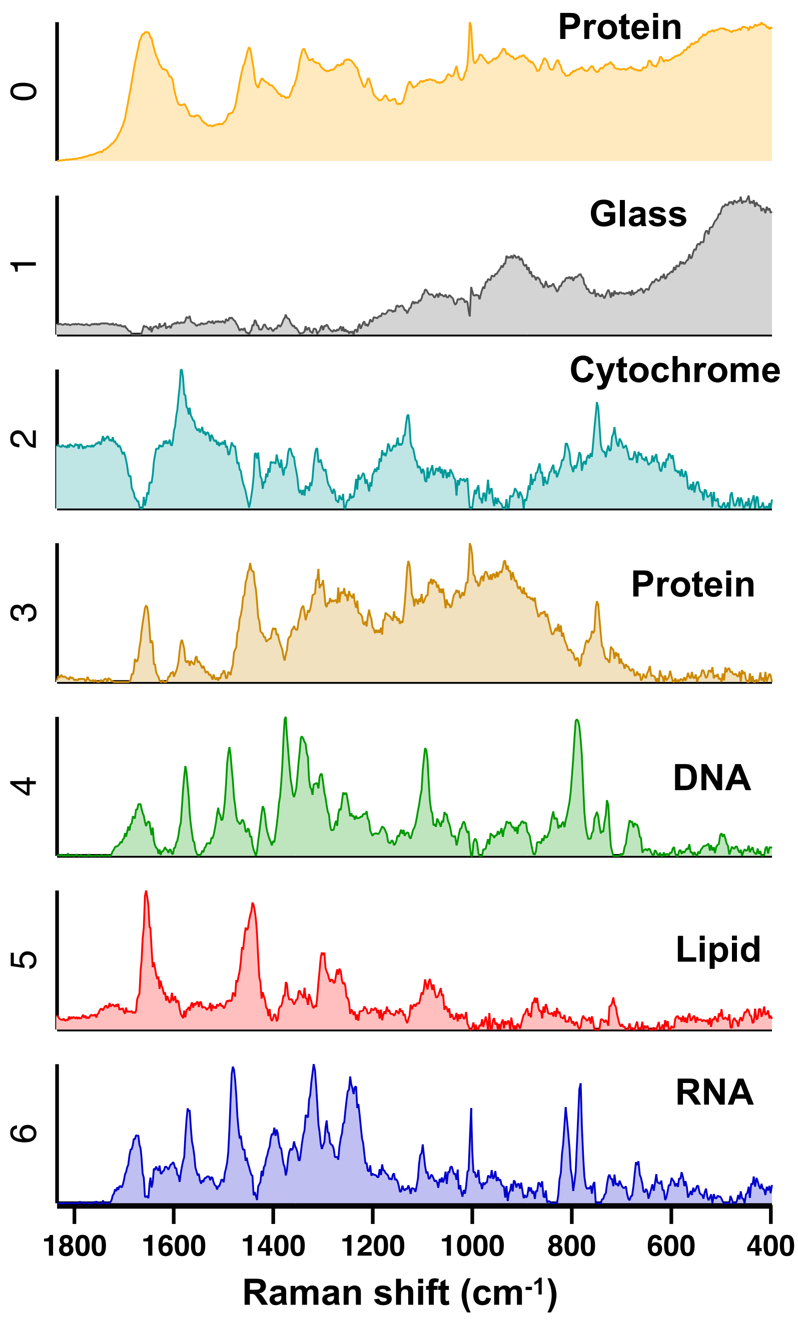


**Supplementary Fig. 14. MCR spectra for nucleus regions exclusively.** MCR spectra of nuclei of **NIH3T3** cells. $n$ = 10 cells. The 5th component spectrum was assigned to lipids.

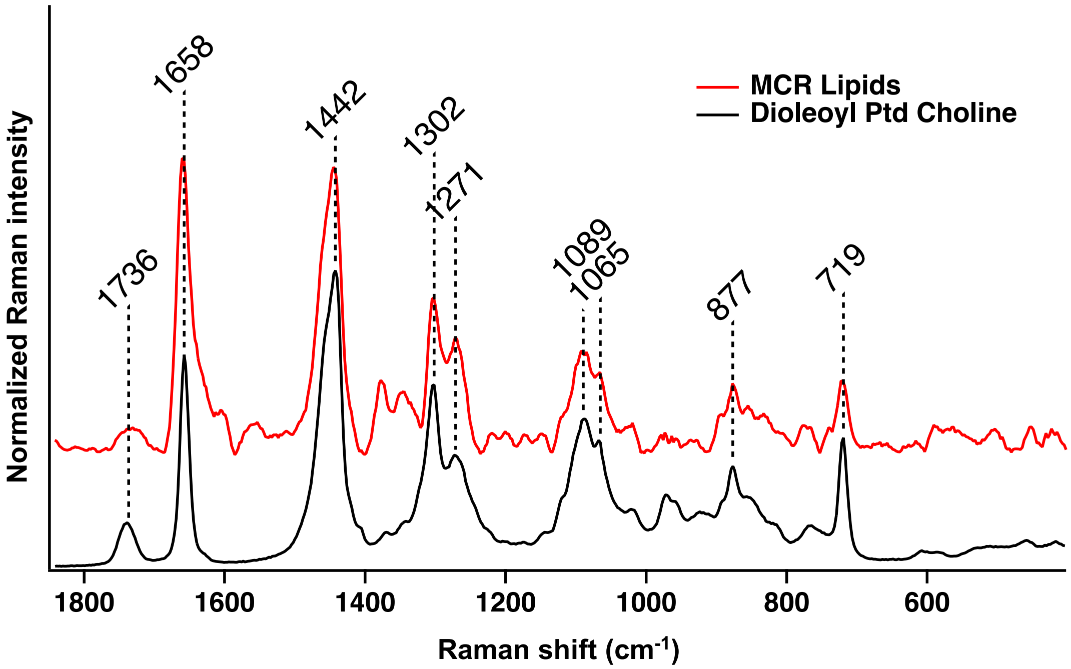


**Supplementary Fig. 15. Phosphatidylcholine is the main component of lipids in the nucleus.** The MCR spectrum assigned to lipids (red) and a Raman spectrum of dioleoyl phosphatidylcholine (DOPC, black). The MCR spectrum exhibits a spectral shape similar to DOPC, indicating that the main component of lipids in the nucleus is a phosphatidylcholine (PC) head group. The MCR spectrum was obtained by analyzing nucleus regions alone and is shown in Supplementary Fig. 14.

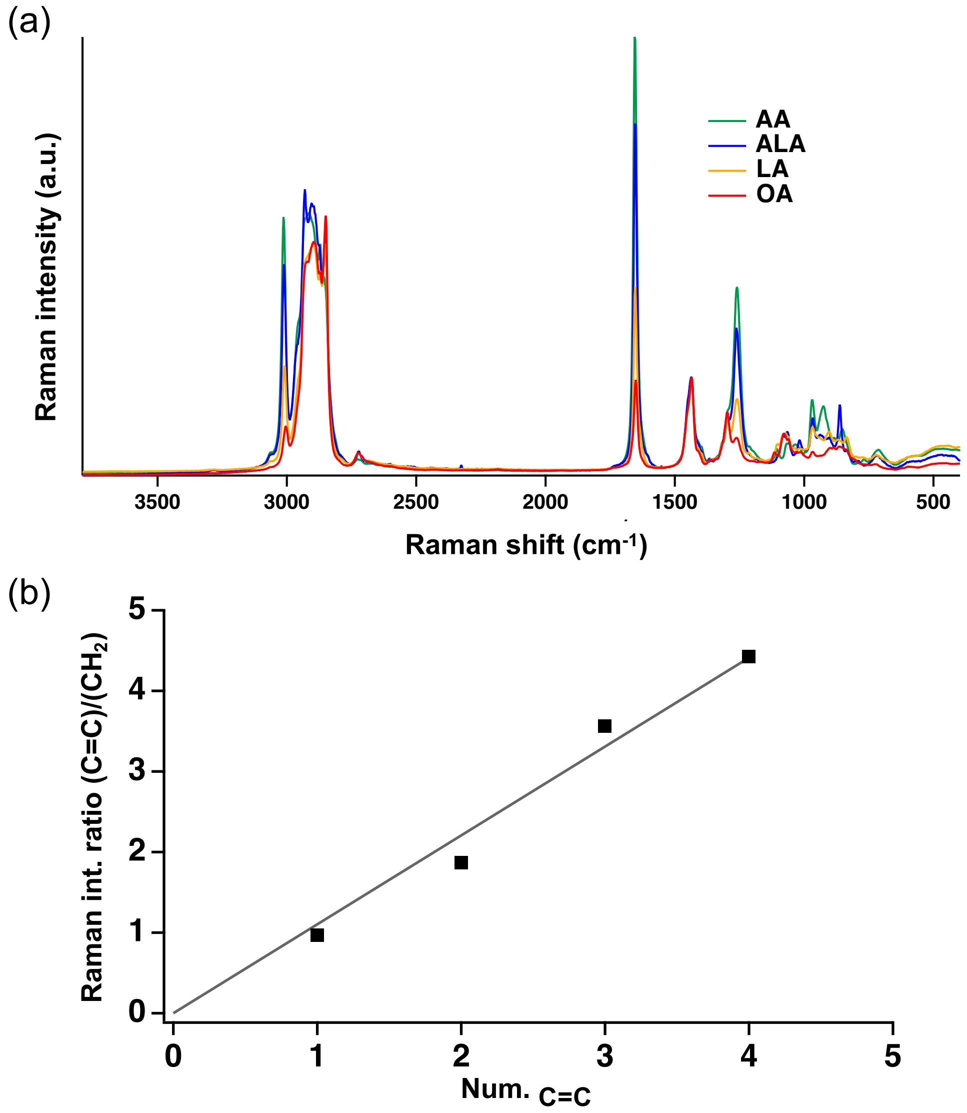


**Supplementary Fig. 16. Raman spectra provide information on lipid unsaturation levels.** (a) Raman spectra of arachidonic acid (AA, green), α-linolenic acid (ALA, blue), linoleic acid (LA, yellow), and oleic acid (OA, red). (b) The intensity ratio of the Raman bands between the C=C stretching band at 1650 $cm^{-1}$ and the $CH_2$ scissors band at 1440 $cm^{-1}$ as a function of the number of C=C bonds. The intensity ratio is proportional to the number of C=C bonds and can be used to evaluate the unsaturation level of lipids.

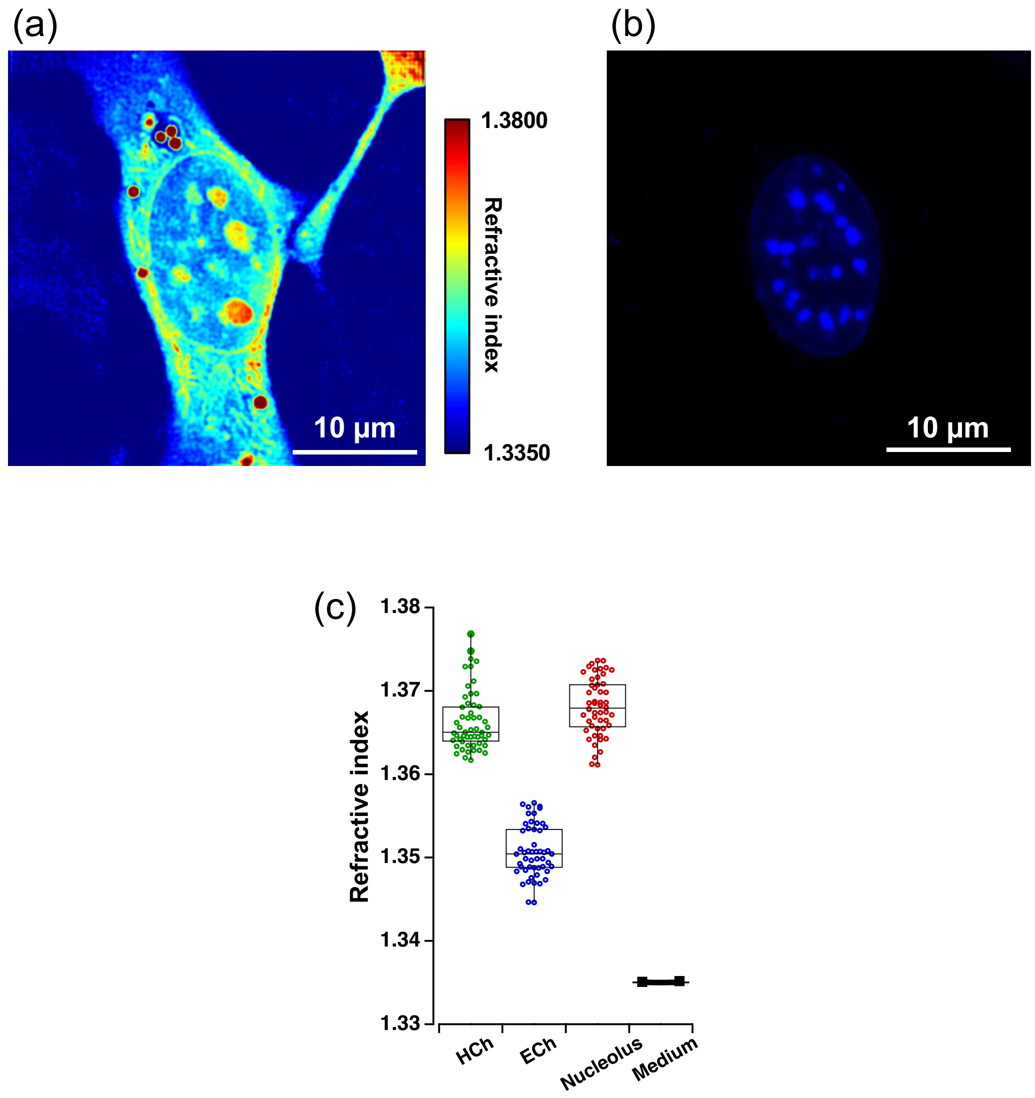


**Supplementary Fig. 17. Refractive index of nuclear compartments.** (a, b) Refractive index (a) and fluorescence (b) images of a Hoechst-stained NIH3T3 cell. (c) Refractive index of each nuclear compartment ($n$ = 18 cells). Refractive index imaging allows visualization of heterochromatin in NIH3T3 cells as regions of high refractive index, even without Hoechst staining.

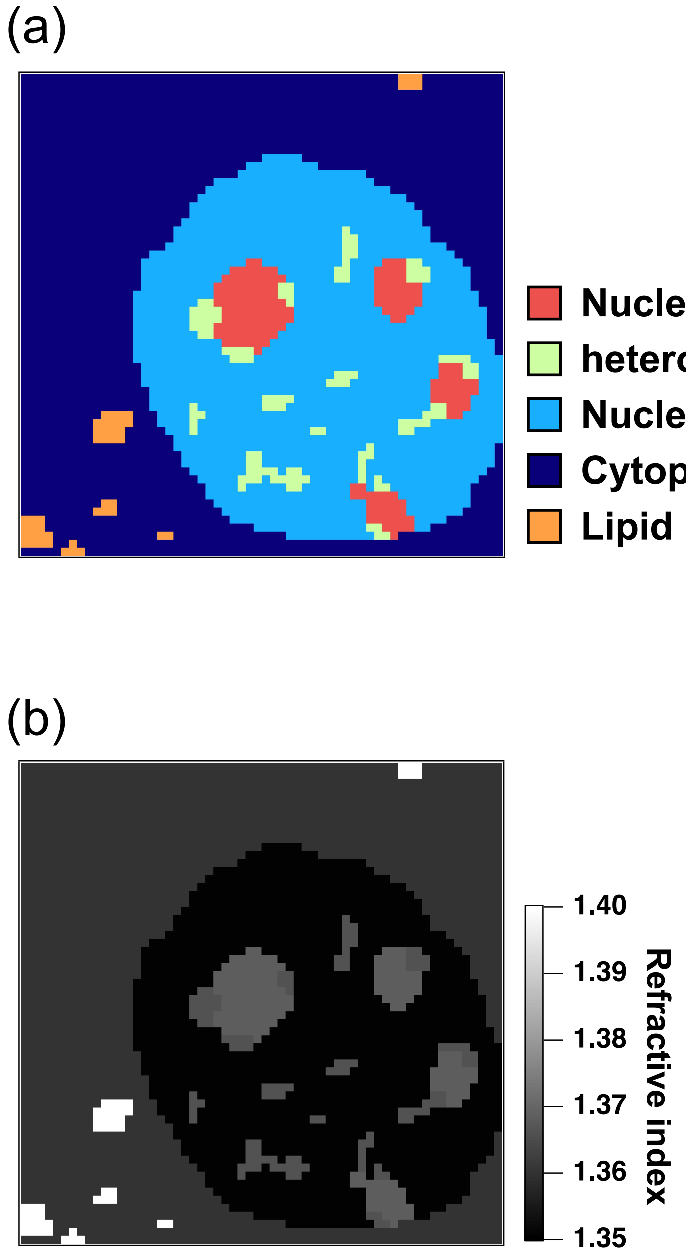


**Supplementary Fig. 18. Refractive index map of an NIH3T3 cell based on Raman imaging.** (a) A color-coded image of cellular compartments based on the MCR images shown in Fig. 2 (in the main text): nucleolus (red), heterochromatin (green), nucleoplasm (light blue), cytoplasm (dark blue), and lipid droplet (orange). (b) A refractive index map generated by assigning the average refractive index value to each cellular compartment.

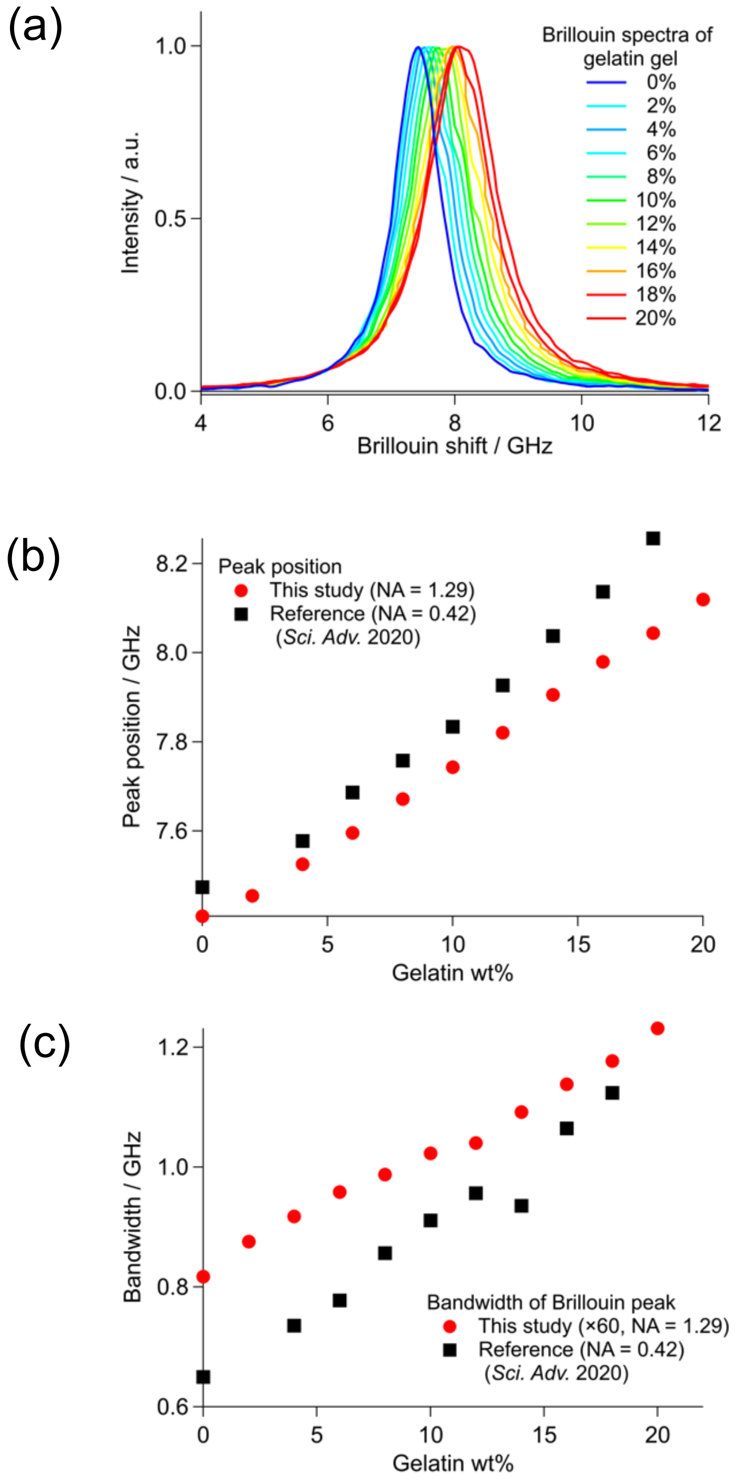


**Supplementary Fig. 19. Effects of high-numerical-aperture (NA) objective lens on Brillouin spectra.** (a) Brillouin spectra of gelatin gels obtained using an objective lens with an NA of 1.23. (b, c) Comparison of the peak position (b) and bandwidth (c) between this study and the previous study using a low NA objective lens[2].

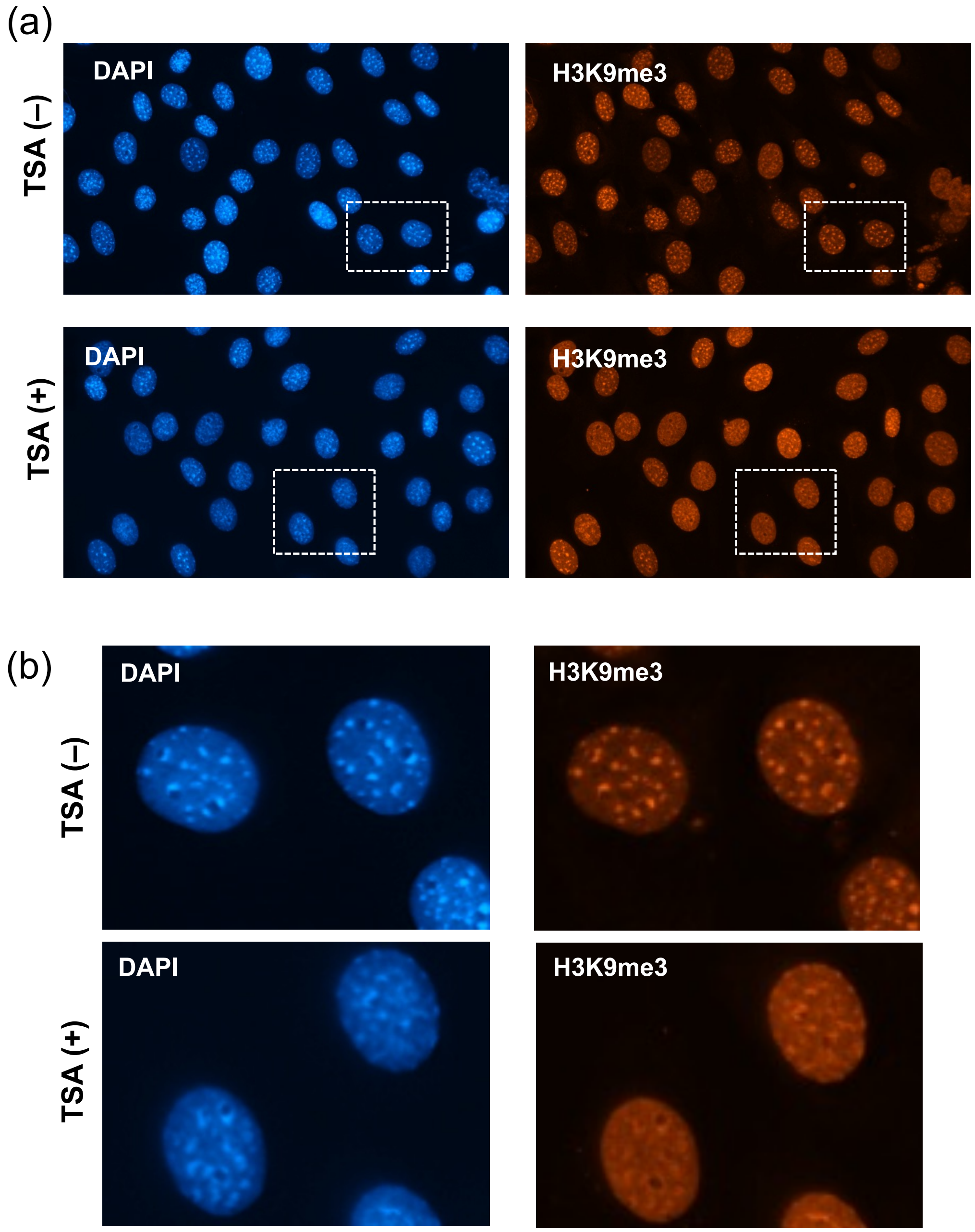


**Supplementary Fig. 20. Chromocenters in trichostatin A (TSA)-treated cells.** (a, b) Large-scale (a) and magnified (b) fluorescence images of NIH3T3 cells stained with DAPI and H3K9me3 antibody before and after TSA treatment. After TSA treatment, heterochromatin appears larger and less distinct, indicating a decrease in the difference between the interior and exterior regions.

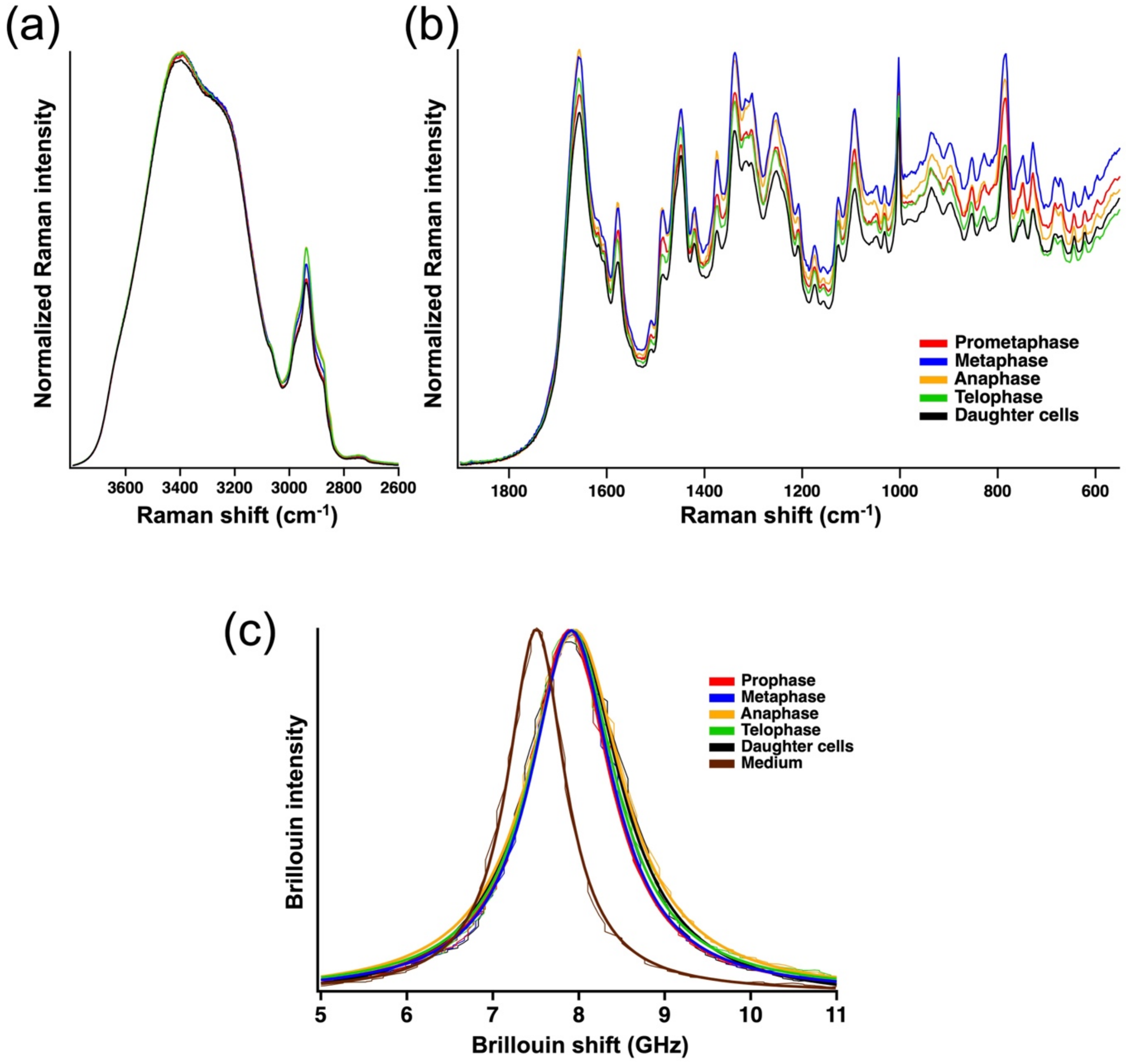


**Supplementary Fig. 21. Raman and Brillouin spectra of NIH3T3 cells in mitosis.** (a, b) Raman spectra of NIH3T3 cells in mitosis and daughter cells in the high-wavenumber region (a) and fingerprint region (b). (c) Brillouin spectra of NIH3T3 cells in mitosis and daughter cells along with the medium spectrum. $n$ = 5 for prometaphase, 5 for metaphase, 7 for anaphase, 5 for telophase, and 6 for daughter cells.

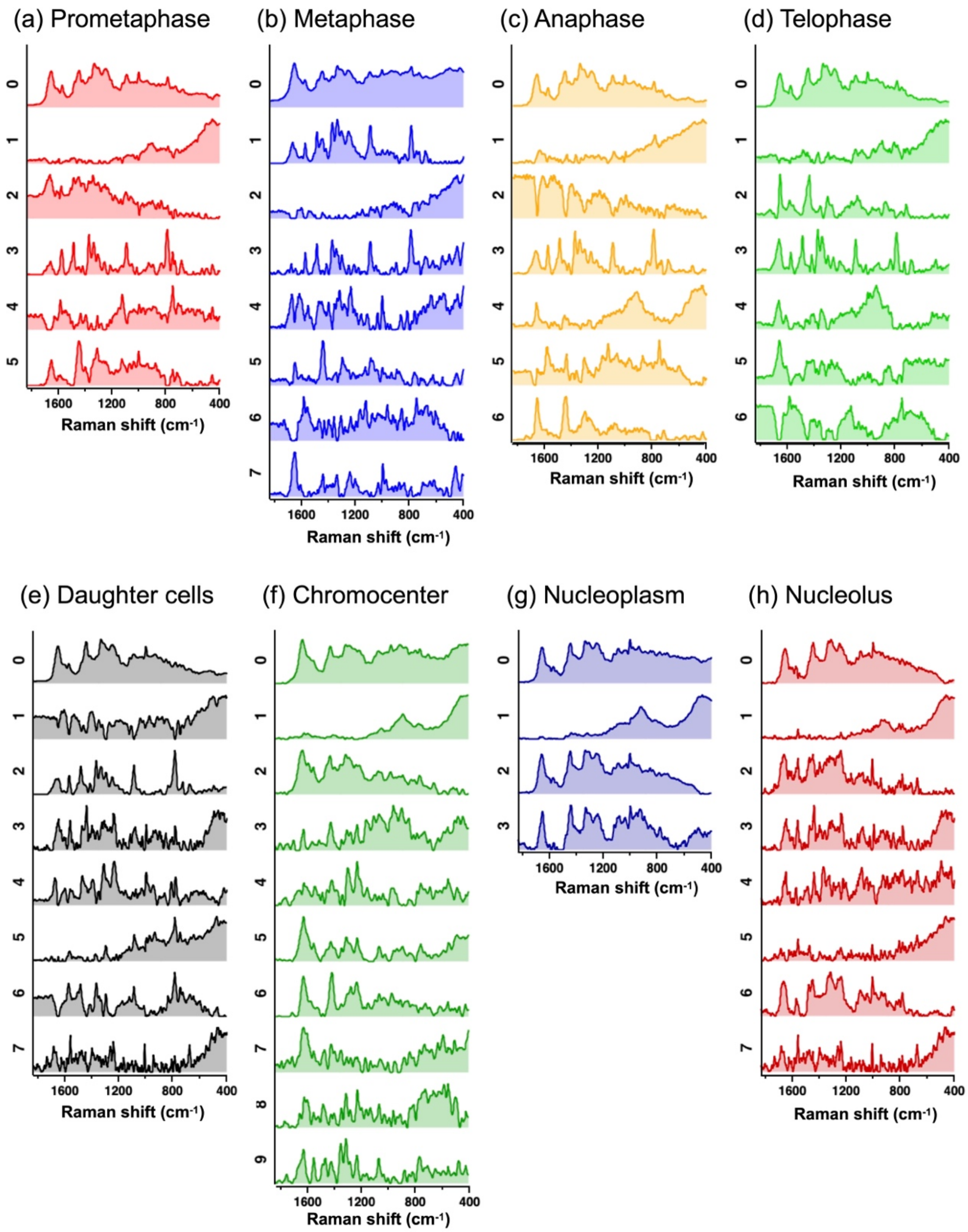


**Supplementary Fig. 22. MCR spectra of mitotic and interphase NIH3T3 cells.** (a-e) Series of MCR component spectra of mitotic cells in prometaphase (a), metaphase (b), anaphase (c), and telophase (d), and their daughter cells (e). (e-g) Series of MCR component spectra derived exclusively from chromocenter (e), nucleoplasm (f), and nucleoli regions of interphase NIH3T3 cells.

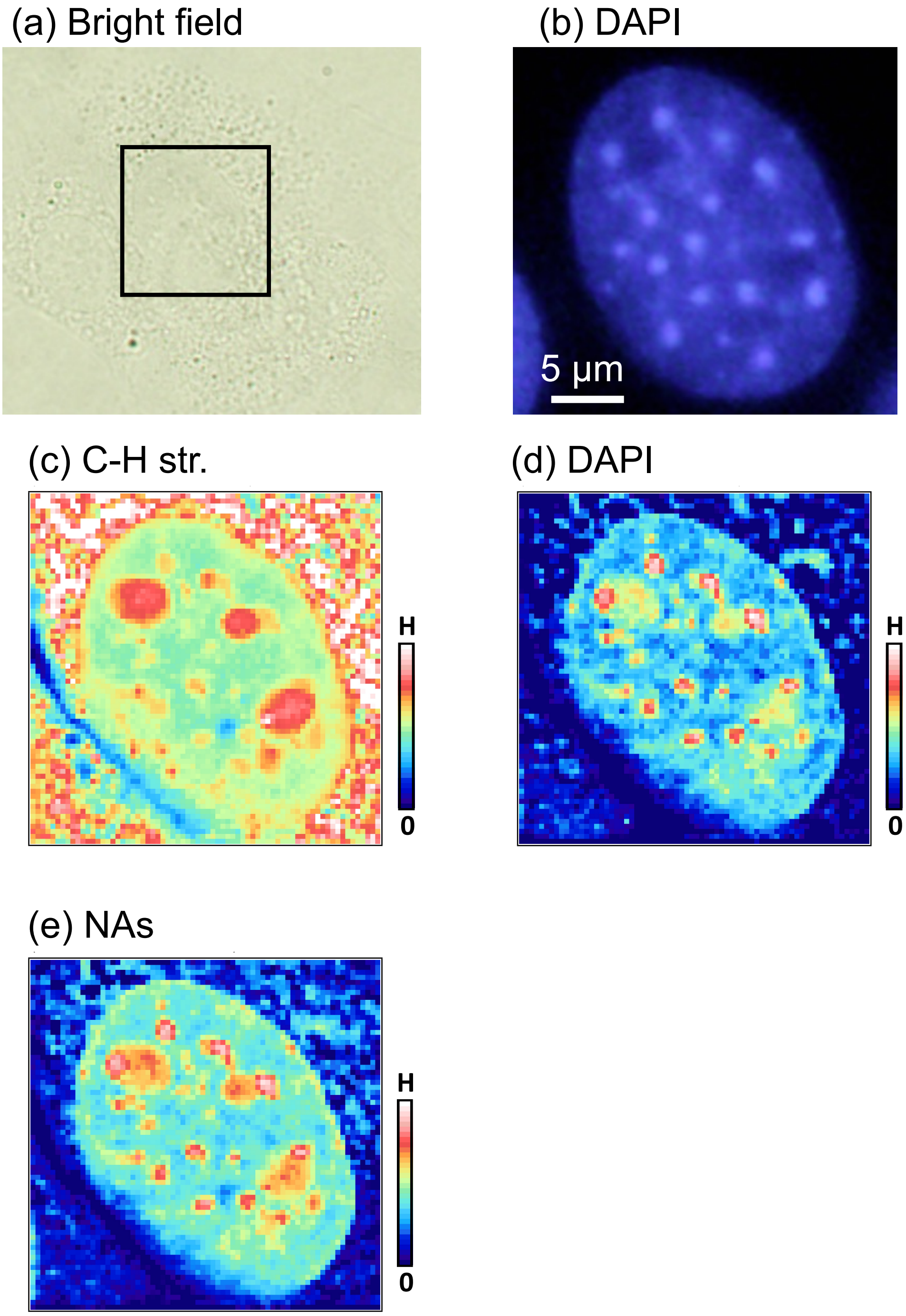


**Supplementary Fig. 23. Raman images of a DAPI-stained NIH3T3 cell.** (a-e) Bright-field (a), DAPI fluorescence (b), and Raman (c-e) images of a fixed and DAPI-stained NIH3T3 cell. Raman images were obtained by mapping the intensity of the C–H stretching region (2800–3000 $cm^{-1}$, c), DAPI resonant band (1590–1630 $cm^{-1}$, d), and the pyrimidine band (780–820 $cm^{-1}$, e).

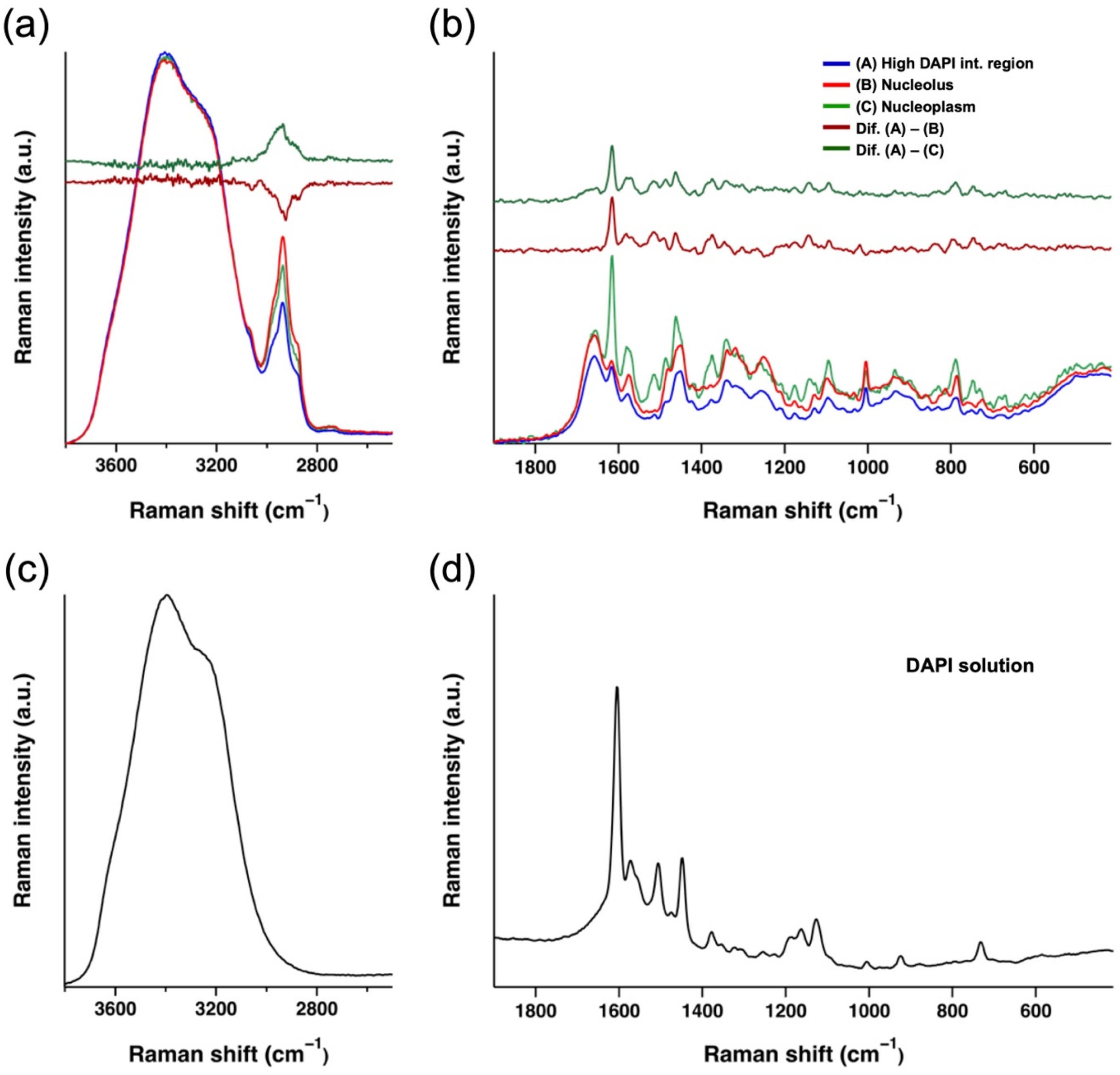


**Supplementary Fig. 24. Raman spectra of DAPI-stained NIH3T3 cells.** (a, b) Raman spectra of the DAPI-intense region (blue), nucleolus (red), and nucleoplasm (green) in DAPI-stained fixed NIH3T3 cells, shown for the high wavenumber (a) and fingerprint (b) regions. (c, d) Raman spectrum of a DAPI solution shown for the high wavenumber (c) and fingerprint (d) regions. DAPI exhibits strong resonant Raman bands around 1450–1650 $cm^{-1}$. Raman bands attributed to lipids, such as the sharp C=C band at 1660 $cm^{-1}$ and the ester band at 1750 $cm^{-1}$, cannot be observed in the difference spectra (high DAPI int. region–nucleolus, high DAPI int. region–nucleoplasm)

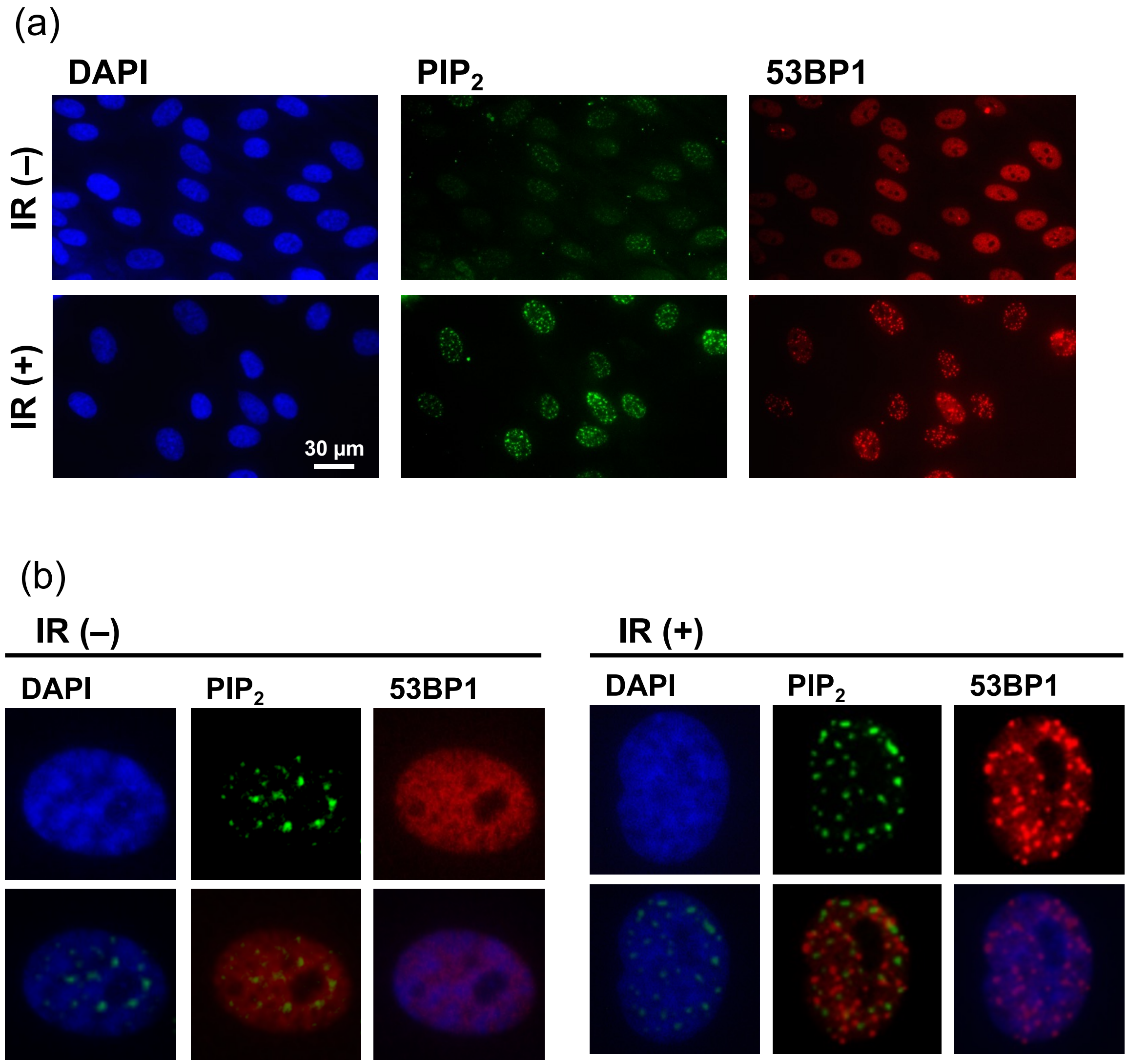


**Supplementary Fig. 25. Immunofluorescence images show that $PIP_2$ does not localize in the chromocenter.** (a, b) Large-scale (a) and magnified (b) immuno-fluorescence images of cells with and without ionizing radiation (IR) showing DAPI (blue), $PIP_2$ (green), and 53BP1 (red) and their merged images. Upon IR exposure, there is an increase in 53BP1-formed foci and the number of $PIP_2$ droplets. However, regardless of IR, DAPI-stained chromocenters do not overlap with 53BP1 or $PIP_2$ droplets.

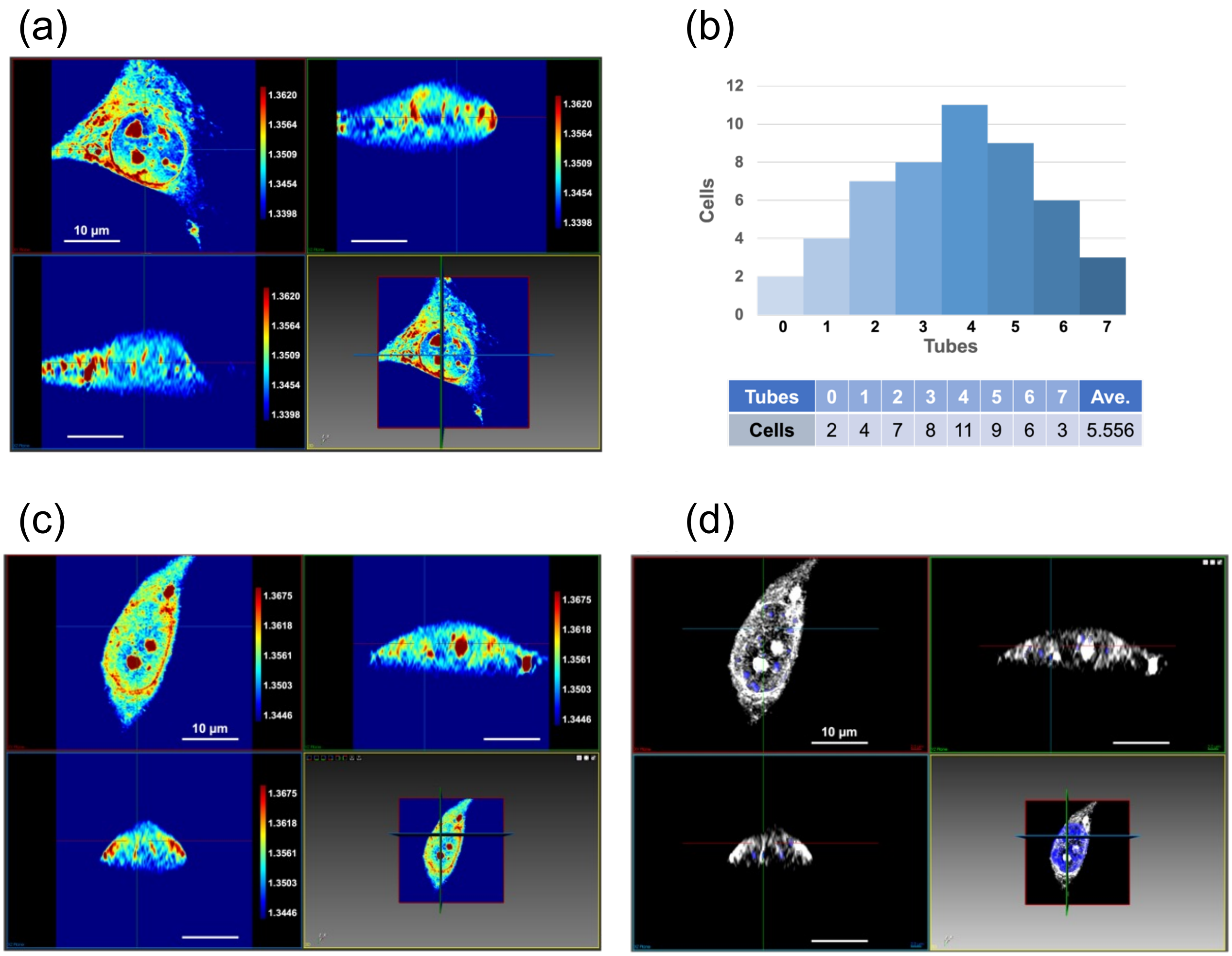


**Supplementary Fig. 26**. **Refractive index images of nuclear membrane invagination tubes.** (a) 3D refractive index images of a NIH3T3 cell. (b) Number distribution of nuclear membrane invagination tubes in NIH3T3 cells in interphase (*n* = 50 cells). (c) 3D refractive index image of a Hoechst-stained NIH3T3 cell. (d) 3D merged image of fluorescence (blue) and refractive index (white). Several tubular structures with high refractive index, attributed to nuclear membrane invagination tubes, were observed within the nucleus.

**Supplementary Table 1.** Assignments of Raman bands[3-10]

| Raman shift ($cm^{-1}$) | Assignment | Compound |
|---|---|---|
| 3015 | =C-H stretch. | Lipids |
| 2940 | $CH_3$ s-stretch. | Proteins/Lipids |
| 2892 | Overtone of $CH_3$ deform. in Fermi resonance with $CH_3$ s-stretch. / $CH_2$ a-stretch. | Proteins/Lipids |
| 2853 | $CH_2$ s-stretch. | Lipids |
| 1748 | C=O stretch. (ester) | Lipids |
| 1658 | cis C=C stretch. / Amide I | Lipids/Proteins |
| 1581 | Purine ring (A and G) | DNAs & RNAs |
| 1573 | Purine ring (A and G) | DNAs & RNAs |
| 1486 | Purine ring (A and G) | DNAs & RNAs |
| 1438 | $CH_2$ scis. | Lipids |
| 1422 | A and G | DNAs & RNAs |
| 1375 | T, A, and C | DNAs |
| 1337 | Purine ring (A and G) | DNAs & RNAs |
| 1323 | CH deform. | Proteins |
| 1307 | $CH_2$ twist. | Lipids |
| 1269 | =C-H bend. | Lipids |
| 1243 | Amide III | Proteins |
| 1212 | C-$C_6H_5$ (phenyl ring) stretch. | Proteins |
| 1092 | $PO_2$ - stretch. | DNAs & RNAs |
| 1005 | Phenyl ring breath. | Proteins |
| 936 | C-C | Proteins |
| 900 | bk. | DNAs & RNAs |
| 818 | bk.(O-P-O) | RNAs |
| 788 | Pyrimidine ring(C, T, and U) / bk.(O-P-O) | DNAs & RNAs |
| 753 | T | DNAs & RNAs |
| 719 | Head group (choline (H3C)N+ sym. stretch.) of phosphatidylcholine | Lipids |
